# SARS-CoV-2 Cell Entry Factors ACE2 and TMPRSS2 are Expressed in the Pancreas but are Not Enriched in Islet Endocrine Cells

**DOI:** 10.1101/2020.08.31.275719

**Authors:** Katie C. Coate, Jeeyeon Cha, Shristi Shrestha, Wenliang Wang, Luciana Mateus Gonçalves, Joana Almaça, Meghan E. Kapp, Maria Fasolino, Ashleigh Morgan, Chunhua Dai, Diane C. Saunders, Rita Bottino, Radhika Aramandla, Regina Jenkins, Roland Stein, Klaus H. Kaestner, Golnaz Vahedi, HPAP consortium, Marcela Brissova, Alvin C. Powers

**Author notes:** Co-first authors (Katie C. Coate, Jeeyeon Cha). Correspondence; Alvin C. Powers or Marcela Brissova, Vanderbilt University Medical Center, 2215 Garland Ave, Nashville, TN 37232-0475.

## Abstract

Reports of new-onset diabetes and diabetic ketoacidosis in individuals with COVID-19 have led to the hypothesis that SARS-CoV-2, the virus that causes COVID-19, is directly cytotoxic to pancreatic islet β cells. This would require binding and entry of SARS-CoV-2 into host β cells via cell surface co-expression of ACE2 and TMPRSS2, the putative receptor and effector protease, respectively. To define ACE2 and TMPRSS2 expression in the human pancreas, we examined six transcriptional datasets from primary human islet cells and assessed protein expression by immunofluorescence in pancreata from donors with and without diabetes. *ACE2* and *TMPRSS2* transcripts were low or undetectable in pancreatic islet endocrine cells as determined by bulk or single cell RNA sequencing, and neither protein was detected in α or β cells from these donors. Instead, ACE2 protein was expressed in the islet and exocrine tissue microvasculature and also found in a subset of pancreatic ducts, whereas TMPRSS2 protein was restricted to ductal cells. The absence of significant ACE2 and TMPRSS2 co-expression in islet endocrine cells reduces the likelihood that SARS-CoV-2 directly infects pancreatic islet β cells through these cell entry proteins.

## Introduction

In coronavirus disease of 2019 (COVID-19), elevated plasma glucose levels with or without pre-existing diabetes and BMI have been identified as independent risk factors of morbidity and mortality with severe acute respiratory syndrome-associated coronavirus-2 (SARS-CoV-2) infection (Barron et al., 2020; Cariou et al., 2020; Holman et al., 2020; Riddle et al., 2020; Wang, 2020; Zhou et al., 2020). Isolated cases of new-onset diabetes and diabetic emergencies such as ketoacidosis and hyperosmolar hyperglycemia have been reported with COVID-19 (Chee et al., 2020; Goldman et al., 2020; Hollstein et al., 2020; Kim et al., 2020; Li et al., 2020a; Rafique and Ahmed, 2020; Unsworth et al., 2020), leading to the hypothesis that SARS-CoV-2 has a diabetogenic effect mediated by direct cytotoxicity to pancreatic islet β cells (Koch, 2020).

*In vitro* studies have shown that SARS-CoV-2 entry into human host cells requires binding to the cell surface receptor angiotensin converting enzyme 2 (ACE2) as well as proteolytic cleavage of the viral spike (S) protein by transmembrane serine protease 2 (TMPRSS2) (Hoffmann et al., 2020; Lan et al., 2020; Shang et al., 2020; Wiersinga et al., 2020). In 2010, Yang and colleagues (Yang et al., 2010) examined autopsy samples from a single deceased patient infected by SARS-CoV-1, which uses similar machinery for binding and cellular entry, and reported expression of ACE2 in pancreatic islet cells. Though the identity of these islet cells was not assessed, the authors suggested that binding of ACE2 by SARS-CoV-1 damages islets and causes acute diabetes, which could be reversed after viral recovery (Yang et al., 2010). There have been occasional reports of other viral infections eliciting a diabetogenic effect (reviewed in (Filippi and von Herrath, 2008)). More recently, Yang and colleagues reported that β-like cells derived from human pluripotent stem cells (hPSCs) as well as β cells of primary human islets express ACE2, raising the possibility of direct infection and cytotoxicity of β cells by SARS-CoV-2 (Yang et al., 2020). Importantly, neither of these prior studies (Yang et al., 2010; Yang et al., 2020) characterized the expression and localization of TMPRSS2, an obligate co-factor for SARS-CoV-2 cellular entry. Thus, a more detailed analysis of both ACE2 and TMPRSS2 expression and localization in human pancreatic tissue from normal donors and those with diabetes is urgently needed. The purpose of this study was to test the hypothesis that native pancreatic islet β cells possess the cellular machinery that could render them direct targets of SARS-CoV-2. Importantly, we found that ACE2 and TMPRSS2 protein are not detectable in human islet endocrine cells from normal donors or those with diabetes, making a direct diabetogenic effect of SARS-CoV-2 via ACE2 and TMPRSS2 unlikely.

## Results and Discussion

### *ACE2* and *TMPRSS2* mRNA expression is minimal in human α or β cells

We first evaluated mRNA expression of *ACE2* and *TMPRSS2* from two existing bulk RNA-sequencing (RNA-seq) datasets (Arda et al., 2016; Blodgett et al., 2015) where human islet α and β cells were enriched by fluorescence activated cell sorting and compared their expression to that of key islet-enriched genes, some of which are normally expressed at relatively low levels in islet cells (e.g., transcription factors). Median expression level of *ACE2* and *TMPRSS*2 mRNA was much less than transcripts of such key islet-enriched genes in human α and β cells (~84% and 92% lower than α and β cell-enriched transcripts, respectively) (**Figure 1A;** n=7-8 adult donors per study). In addition, analysis of four single-cell (sc) RNA-seq datasets of human pancreatic cells (Baron et al., 2016; Camunas-Soler et al., 2020; Kaestner et al., 2019; Segerstolpe et al., 2016) revealed that, in aggregate, less than 1.5% of β cells expressed *ACE2* or *TMPRSS2*, and each transcript was minimally expressed or undetectable in all other endocrine cell subsets (**Figure 1B, S4J** and **Table S1**). We note that this includes analysis of the robust Human Pancreas Analysis Program (HPAP) dataset that includes more than 25,000 cells from 11 normal donors (Kaestner et al., 2019), findings of which are confirmed in the analyses of three previously reported, but smaller, datasets (**Figure 1B** and **S4J**). Furthermore, given that *ACE2* and *TMPRSS2* co-expression is required for canonical SARS-CoV-2 host cell entry (Hoffmann et al., 2020), we also evaluated this occurrence but found that no β cells co-expressed *ACE2* and *TMPRSS2* in any of these four datasets (**Table S1**).

**Figure 1.**
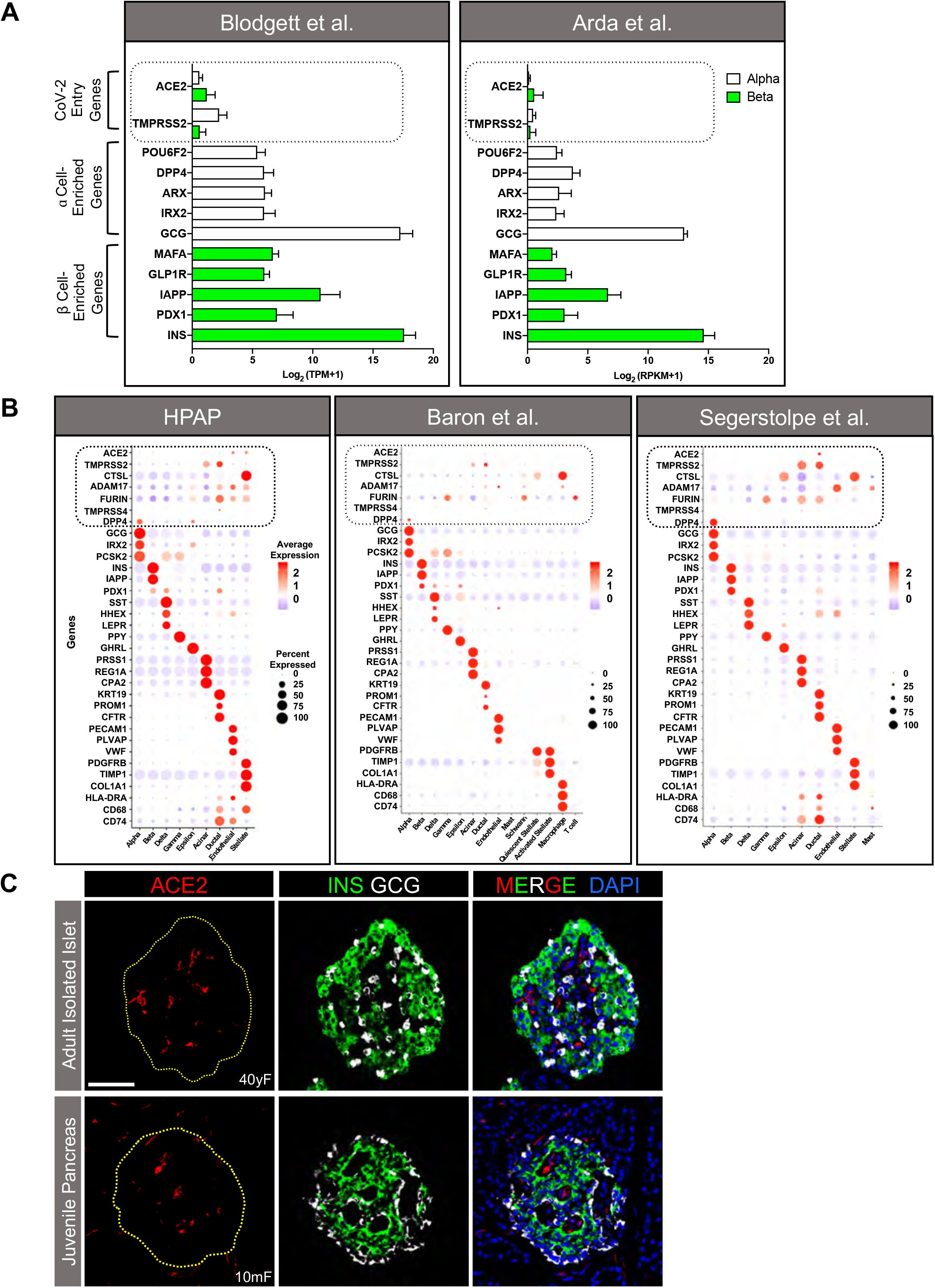
*ACE2* and *TMPRSS2* Expression is Minimal in Isolated Human Islet α and β Cells. (**A**) Relative expression of *ACE2* and *TMPRSS2* compared with select α (white bars) and β (green bars) cell type-enriched genes in sorted human islet α and β cells from previously published bulk RNA-seq datasets, reported as transcript per million mapped reads (TPM; n=7; (Blodgett et al., 2015)) or reads per kilobase of transcript per million mapped reads (RPKM; n=8; (Arda et al., 2016)). Mean expression values are presented as log2 (TPM+1) or log2 (RPKM+1) to account for negative values. Dotted line highlights *ACE2* and *TMPRSS2* expression. (**B**) Dot plots of *ACE2*, *TMPRSS2*, *CTSL*, *ADAM17*, *FURIN*, *TMPRSS4*, and *DPP4* expression compared with cell type-enriched genes from three single cell (sc) RNA-seq datasets (Baron et al., 2016; Kaestner et al., 2019; Segerstolpe et al., 2016). Dot size indicates percentage of cells in a given population expressing the gene; dot color represents scaled average expression. Dotted lines highlight expression of the putative SARS-CoV-2 entry machinery. Percentages of β cells expressing and co-expressing these genes are available in Table S1. (**C**) Representative images showing an isolated islet from an adult human donor (top panels) and a pancreatic section from a juvenile human donor (bottom panels) stained for ACE2 (red; antibody ab15348), insulin (INS; green), glucagon (GCG; white), and DAPI (blue). Dotted yellow line denotes islet area. Scale bar is 50 μm. Human pancreatic donor information is available in Table S2 (1C, donors I1-I3, J1-J5). See also Figures S1, S2, and S4.

As for non-endocrine cells, the HPAP dataset (Kaestner et al., 2019) revealed that a small subset (< 5%) of endothelial and stellate cells (which include pericytes) expressed moderate to high levels of *ACE2*, whereas only ~1-3% of either population expressed *ACE2* in the datasets by Baron and colleagues (Baron et al., 2016) and Segerstolpe and colleagues (Segerstolpe et al., 2016) (**Figure 1B**). This difference most likely stems from the number of cells analyzed in that the HPAP dataset contains ~2.5- and 20-fold more cells than that of Baron et al. (Baron et al., 2016) or Segerstolpe et al. (Segerstolpe et al., 2016), respectively. In addition, pooled analysis of these datasets (Baron et al., 2016; Camunas-Soler et al., 2020; Kaestner et al., 2019) showed that less than 1% of acinar or ductal cells expressed *ACE2*, whereas ~35% of both cell types expressed *TMPRSS2*. In the dataset by Segerstolpe and colleagues (Segerstolpe et al., 2016), *ACE2* was expressed in ~20% of acinar and ductal cells, whereas *TMPRSS2* was expressed in greater than 75% of both populations (**Figure 1B**). Evaluation of *ACE2* and *TMPRSS2* co-expression in non-endocrine cells revealed that on average, less than 1% of acinar, ductal, endothelial and stellate cells co-expressed both transcripts in these datasets (Baron et al., 2016; Camunas-Soler et al., 2020; Kaestner et al., 2019). In the Segerstolpe et al. dataset (Segerstolpe et al., 2016), ~5% of acinar cells and 15% of ductal cells co-expressed *ACE2* and *TMPRSS2*, but this difference could represent unintended cell selection bias as this study analyzed the fewest number of cells.

Altogether, these gene expression findings reduce the likelihood that SARS-CoV-2 can bind and enter human β cells via the canonical pathway involving ACE2 and TMPRSS2. However, they do not exclude the possibility of viral entry via non-canonical pathways involving other suggested effector proteases of SARS-CoV-2, such as Cathepsin L (*CTSL*), ADAM metallopeptidase domain 17 (*ADAM17*), *FURIN* and *TMPRSS4* (Breidenbach et al., 2020; Schreiber et al., 2020; Seyedpour et al., 2020; Zang et al., 2020). We evaluated the expression of these transcripts across all four scRNA-seq datasets and found that with the exception of *TMPRSS4*, each is ubiquitously expressed to varying degrees in all cell types (**Figure 1B, Table S1**). In aggregate, however, less than 1% of β cells co-expressed *ACE2* with *CTSL*, *ADAM17*, *FURIN* or *TMPRSS4* (**Table S1**). An outstanding question is whether expression of these transcripts in even 1% of β cells is sufficient to confer permissiveness to SARS-CoV-2 infection *in vivo*. Furthermore, heparan sulfate in the extracellular glycocalyx (Clausen et al., 2020) and motile cilia (present on islet endocrine cells) (Lee et al., 2020) were mechanisms proposed to assist ACE2-mediated viral entry. Future studies addressing these possibilities in β cells are urgently needed. Notwithstanding, *CTSL*, *ADAM17*, *FURIN* and *TMPRSS4* were more highly expressed in exocrine, endothelial and stellate cells, as was ACE2 (**Figure 1B**). These gene expression patterns raise the possibility that infection of certain pancreatic cell types by SARS-CoV-2 indirectly impacts β cell function.

Obesity is a key metabolic risk factor associated with mortality with COVID-19. A recent report noted increased correlation of *TMPRSS2*, but not *ACE2*, expression with BMI (Taneera et al., 2020). We evaluated the expression of *ACE2*, *TMPRSS2* and *ADAM17* in β cells from 11 donors according to their BMI in our largest sc-RNAseq dataset (Kaestner et al., 2019). We did not observe changes in the expression of *ACE2* or *TMPRSS2* expression with increasing BMI but observed a trend towards higher *ADAM17* expression (**Figure S1**). However, this dataset included only one patient with an obese BMI. Studies with a larger number of donor islets are needed to address the role of obesity and other pancreas-resident SARS-CoV-2 effector molecules on β cell function in the context of COVID-19.

### ACE2 and TMPRSS2 protein are not detected in adult or juvenile human islet α or β cells

Recently, Yang and colleagues (Yang et al., 2020) reported that ACE2 protein was present in α and β cells from primary human islets and in β-like cells derived from hPSCs. To investigate ACE2 expression, we immunostained cryosections of intact isolated human islets using the same ACE2 antibody as used by Yang and colleagues (Yang et al., 2020) (antibody validation studies in **Figure S2**). In accordance with the transcriptomic data reported above, ACE2 was not detected in insulin-positive β cells or glucagon-positive α cells in our samples; instead, ACE2 signal was prominent in surrounding cells such as those of the microvasculature (**Figure 1C, top panels**).

Since hPSC-derived β-like cells differentiated *in vitro* are considered “juvenile-like” and not functionally mature β cells (Nair et al., 2019), one possibility is that ACE2 is expressed in less differentiated cells, or in the cultured, immortalized β cells (EndoCβH1) studied by Fignani et al. (preprint: Fignani, 2020). We investigated such a possibility by staining for ACE2 in pancreatic sections from juvenile human donors (age range 5 days to 5 years) using the same ACE2 antibody as used by Yang et al. (Yang et al., 2020). However, ACE2 was not detected in these juvenile β or α cells (**Figure 1C, bottom panels**), making this explanation less likely.

To further characterize ACE2 and TMPRSS2 protein expression in the native pancreas, we next analyzed human pancreatic tissue sections from normal donors (ND, n=14; age range 18-59 years) and those with type 2 diabetes (T2D, n=12; age range 42-66 years) or type 1 diabetes (T1D, n=11; age range 13-63 years) for these proteins on the same sections. ACE2 protein did not co-localize with markers of α or β cells but was detected within the distinct islet areas indicative of microvascular structures in all ND individuals (**Figures 2A-C”, 2D-F”** and **S3A-D**) as well as those with T2D (**Figures 2G-I”, 2J-L”** and **S3I-L**) or T1D (**Figures 2M-O”, 2P-R”** and **S3O-R**). We noted similar findings with three additional ACE2-directed antibodies, including the same antibodies and dilutions reported by Yang et al. (Yang et al., 2020) and by Fignani et al. (preprint: Fignani, 2020), and one antibody validated by the Human Protein Atlas (HPA) (Hikmet et al., 2020) (**Figure S2**). Three of these antibodies detect epitopes corresponding to the extracellular domain of ACE2 (AF933; MAB933; HPA000288), while one detects an epitope in the C-terminal domain (ab15348). Staining patterns were similar across all antibodies (**Figure S2**). Furthermore, ACE2 antibody specificity (ab15348) was confirmed via peptide competition with the commercially available immunizing peptide (**Figure S2F-I**).

**Figure 2.**
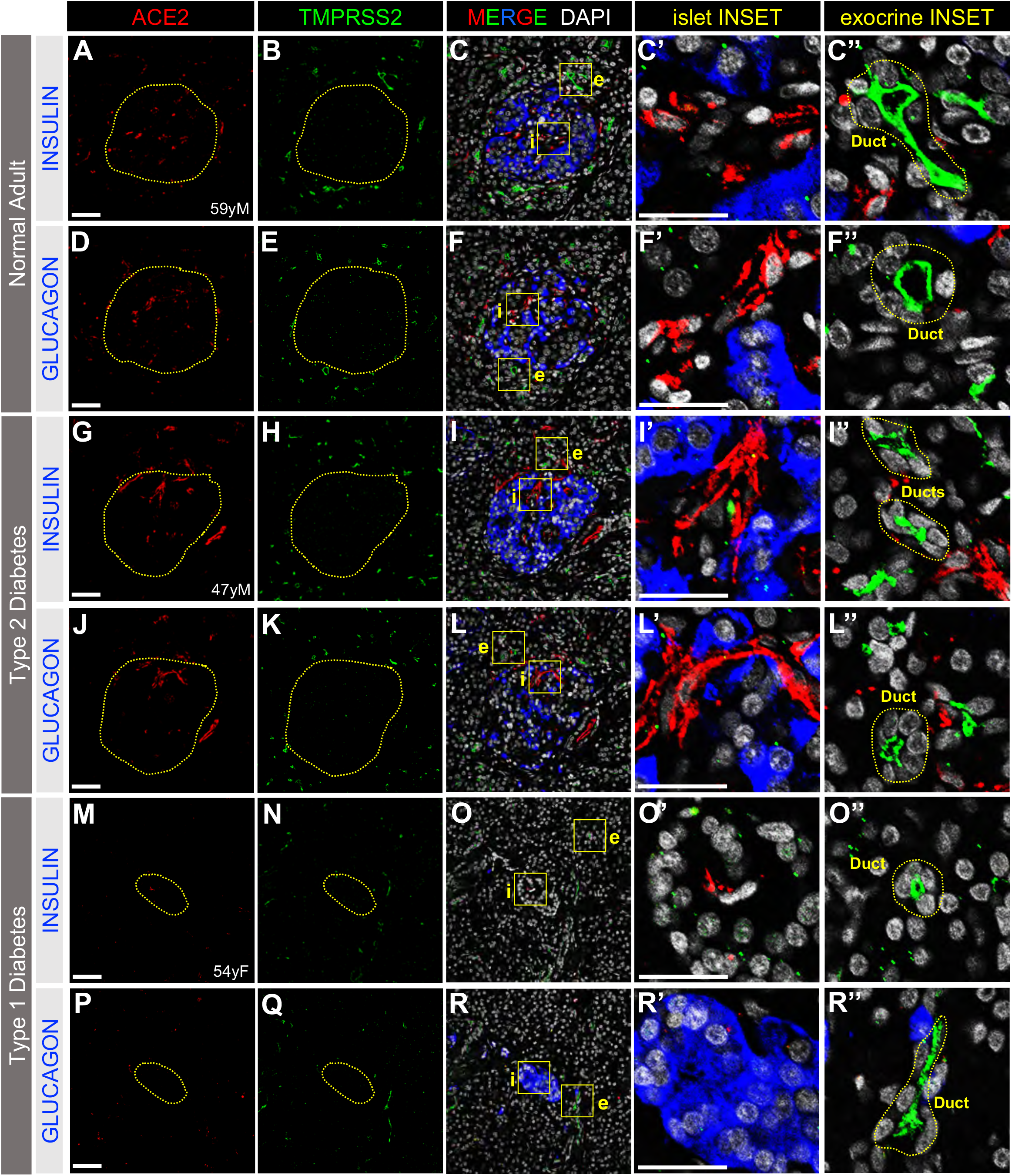
ACE2 and TMPRSS2 Protein are Not Detected in Human Islet α or β Cells in Adult Pancreas. Immunostaining does not detect SARS-CoV-2 cell entry markers ACE2 (**A**, **D**, **G**, **J**, **M**, **P**; antibody AF933; in red) or TMPRSS2 (**B**, **E**, **H**, **K**, **N**, **Q;** in green), in islet β cells (INS, blue; **C-C’**, **I-I’**, **O-O’**) or islet α cells (GCG; blue; **F-F’**, **L-L’**, **R-R’**) in native pancreatic sections from adult donors without diabetes (**A-F”**) or from donors with type 2 (**G-L”**) or type 1 (**M-R”**) diabetes. Dotted yellow lines denote islets. Islet (i) and exocrine (e) inset areas are marked by yellow boxes in MERGE column with DAPI counterstain. Pancreatic ducts, identified structurally by rosette pattern of DAPI-labeled nuclei (white), are shown within dotted yellow lines in exocrine INSET column. Scale bars are 50 μm (**A-R**) and 25 μm (Insets, **C’-R”**). Human pancreatic donor information is available in Table S2 (**A-F”**, donor N9; **G-L”**, donor 2H; **M-R”**, donor 1C). See also Figures S2, S3, and S4.

TMPRSS2 protein was not detected within islets in ND (**Figures 2A-C”, 2D-F”** and **S3E-H**), T2D (**Figures 2G-I”, 2J-L”** and **S3M-N**), or T1D donors (**Figures 2M-O”, 2P-R”** and **S3S-V**). The TMPRSS2-positive structures in exocrine tissue resembled intercalated and larger ducts as indicated by the typical organization of their nuclei (**Figures 2C”, 2F”, 2I”, 2L”, 2O” and 2R”**). We did not detect differences in the signal intensity or spatial distribution of ACE2 or TMPRSS2 between ND and T2D tissue (**Figures 2** and **S3**) but labeling for both proteins appeared to be reduced in T1D pancreatic sections (**Figure 2M-R” and S3O-V**). Altogether, these observations indicate that the cell surface proteins required for canonical SARS-CoV-2 host cell entry were not detected on islet endocrine cells from normal or diabetic donors.

Recent in silico analyses by Vankadari et al. (Vankadari and Wilce, 2020) and Li et al. (Li et al., 2020b) suggested that human dipeptidyl peptidase 4 (DPP4) may interact with SARS-CoV-2 and facilitate its entry into host cells. Therefore, we examined the distribution of DPP4 in human pancreas and found that it localized to α cells, but not β cells, of ND, T2D and T1D islets (**Figure S4A-I**). This finding is consistent with our analysis of four scRNA-seq datasets showing *DPP4*-enriched α cells (**Figures 1B and S4J**). Given that we did not detect TMPRSS2 within islets, these data suggest that DPP4 is an unlikely mediator of β cell entry by SARS-CoV-2 (Drucker, 2020).

The discrepancy between our findings and those of recent studies identifying ACE2 and TMPRSS2 in human β cells (Yang et al., 2010) (Yang et al., 2020) (preprint: Fignani, 2020) is likely explained by important differences in experimental approaches and contexts. Yang et al. (Yang et al., 2010) showed ACE2 staining in a single donor islet but did not identify its cellular identity with endocrine markers. Furthermore, the ACE2 antibody used in their study was not reported, which precluded replication of their finding. Fignani et al. (preprint: Fignani, 2020) identified ACE2 protein in presumed islet endocrine cells from seven non-diabetic donors; however, ACE2 was primarily seen in subcellular compartments rather than the expected cell surface location. Colocalization of ACE2 with insulin granules in this study raises the possibility of a staining artifact. Furthermore, Yang et al. (Yang et al., 2020) examined ACE2 expression in isolated dispersed human islet cells, whereas our study examined ACE2 in isolated intact islets and in human islets of native pancreata. It is possible that dispersion of pancreatic cells may have altered the expression of ACE2 in this context (van den Brink et al., 2017). Likewise, sources of variability in immunostaining techniques can stem from differences in tissue fixation procedures, antigen retrieval methods, antibodies used and their dilutions. Notwithstanding, the remarkable concordance of our findings with those from a recent independent effort in native pancreas (Kusmartseva, et al) suggest that differences in experimental conditions may explain the discordance with other studies.

### ACE2 is localized to islet and exocrine tissue capillaries

The non-endocrine cell staining pattern of ACE2 in islets prompted us to examine whether ACE2 was expressed in the microvasculature, as described in other organs (Hamming et al., 2004). Indeed, staining of adult and juvenile human pancreatic tissue sections with CD31, an endothelial cell marker, revealed that ACE2 labeling was localized to the perivascular compartment of islet capillaries (**Figures 3A-D’, I-L’** and **S5A-H’**). In addition, exocrine tissue capillaries, similar to those in islets, showed perivascular ACE2 labeling (**Figures 3E-H’** and **S5I-P’**). As ACE2-positive cells were scarce in T1D pancreatic tissues (**Figures 2M, 2O’-O”, 2P, 2R’-R”**, and **S3O-R**), the presence of ACE2 labeling in the perivascular compartment was relatively rare in both islet (**Figure S5E-H’**) and exocrine (**Figure S5M-P’**) T1D tissues. The structure and close relationship between ACE2-positive perivascular cells and CD31-positive endothelial cells in islet and exocrine tissue capillaries raised the possibility that ACE2-positive cells were in fact pericytes enveloping capillary endothelial cells of the vascular tube (Almaca et al., 2018). In addition, we observed ACE2-positive cytoplasmic processes extending along CD31-positive endothelial cells colocalized with the extracellular matrix marker collagen-IV within the vascular basement membrane (**Figures 3D’, 3H’**, **S5D’**, **S5H’**, **S5L’ and S5P’**), a pattern that is consistent with pericyte morphology (Almaca et al., 2018).

**Figure 3.**
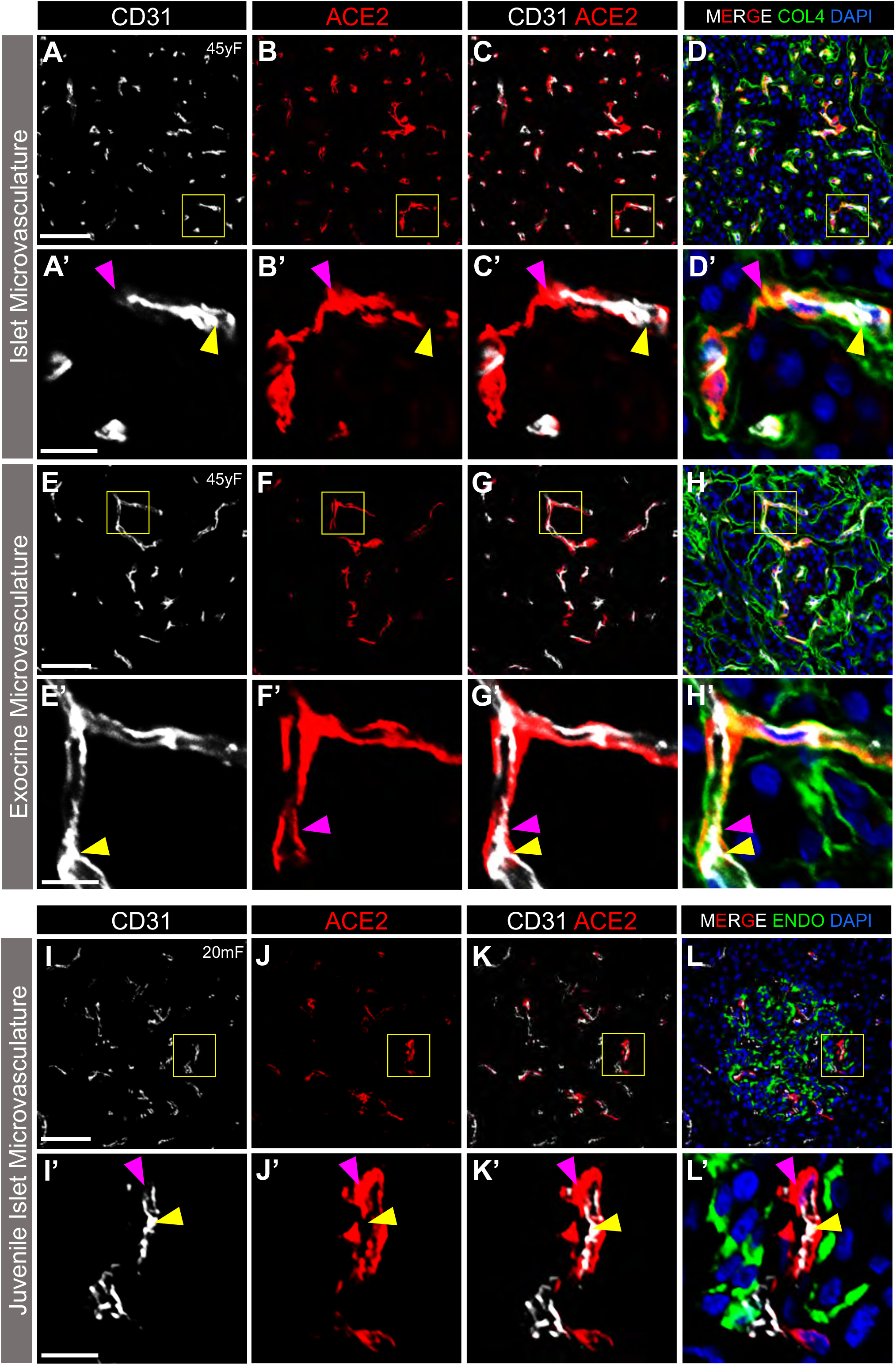
ACE2 Protein is Localized to Islet and Exocrine Capillaries in Adult and Juvenile Human Pancreata. Representative images of endothelial cells (CD31, white) and ACE2-positive perivascular cells (red; antibody ab15348) in the adult islet (**A-D**’) and exocrine (**E-H’**) microvasculature, as well as in juvenile human islet (**I-L’**) microvasculature. Yellow arrowheads point to CD31-positive endothelial cells, while magenta arrowheads point to perivascular ACE2-positive cells. ACE2-positive perivascular cells abut the extracellular matrix marker collagen-IV within the vascular basement membrane (collagen-IV [COL4], green; **D, D’, H, H’**). Inset areas (**A’-L’**) are marked by yellow boxes in **A-L**. In juvenile human pancreas (**I-L’**), ACE2 is not detected in islet endocrine cells expressing insulin and glucagon (ENDO, green); DAPI (blue). Scale bars are 50 μm (**A-L**) and 10 μm (Insets, **A’-L’**). Human pancreatic donor information is available in Table S2 (**A-H’**, donor N2; **I-L’**: donor J4). See also Figure S5.

To address this observation more directly, we visualized ACE2 expression in relationship to known pericyte markers and found that ACE2 was enriched in pericyte populations expressing PDGFRβ (**Figure 4A-D**). Indeed, colocalization analysis revealed that ~60% of ACE2-positive cells were PDGFRβ-positive (**Figure 4H-I**). Since PDGFRβ labels pericytes and perivascular fibroblasts (Almaca et al., 2020), we also stained tissues with a more specific pericyte marker, NG2, and found that ~30% of ACE2-positive cells were NG2-positive (**Figure 4E-G’, 4H-I**). These data agree with our scRNA-seq analysis of the HPAP dataset which identified a subset of pericytes, marked by *PDGFRB*, that express moderate levels of *ACE2* (**Figure 1B**). In addition, our findings support recent reports of pericyte-specific vascular expression of ACE2 in brain and heart tissue (Chen et al., 2020; He, 2020). Emerging evidence from both pre-clinical models of SARS-CoV-2 infection (Aid, 2020) and patients with COVID-19 indicate that cells of the microvasculature (e.g. endothelial cells and pericytes) are requisite contributors to the initiation and propagation of severe disease (Teuwen et al., 2020). Although we found that *TMPRSS2* expression was negligible in endothelial and stellate cells (which includes pericytes) of human islets, *CTSL*, *ADAM17*, *FURIN* and *ACE2* were moderately to highly expressed in these populations (**Figure 1B**), raising the intriguing possibility of direct infection of cells in the islet microvasculature by SARS-CoV-2. Such an occurrence could trigger β cell dysfunction and the metabolic sequelae of COVID-19. Future studies addressing this possibility in human islets and pancreatic tissue are urgently needed.

**Figure 4.**
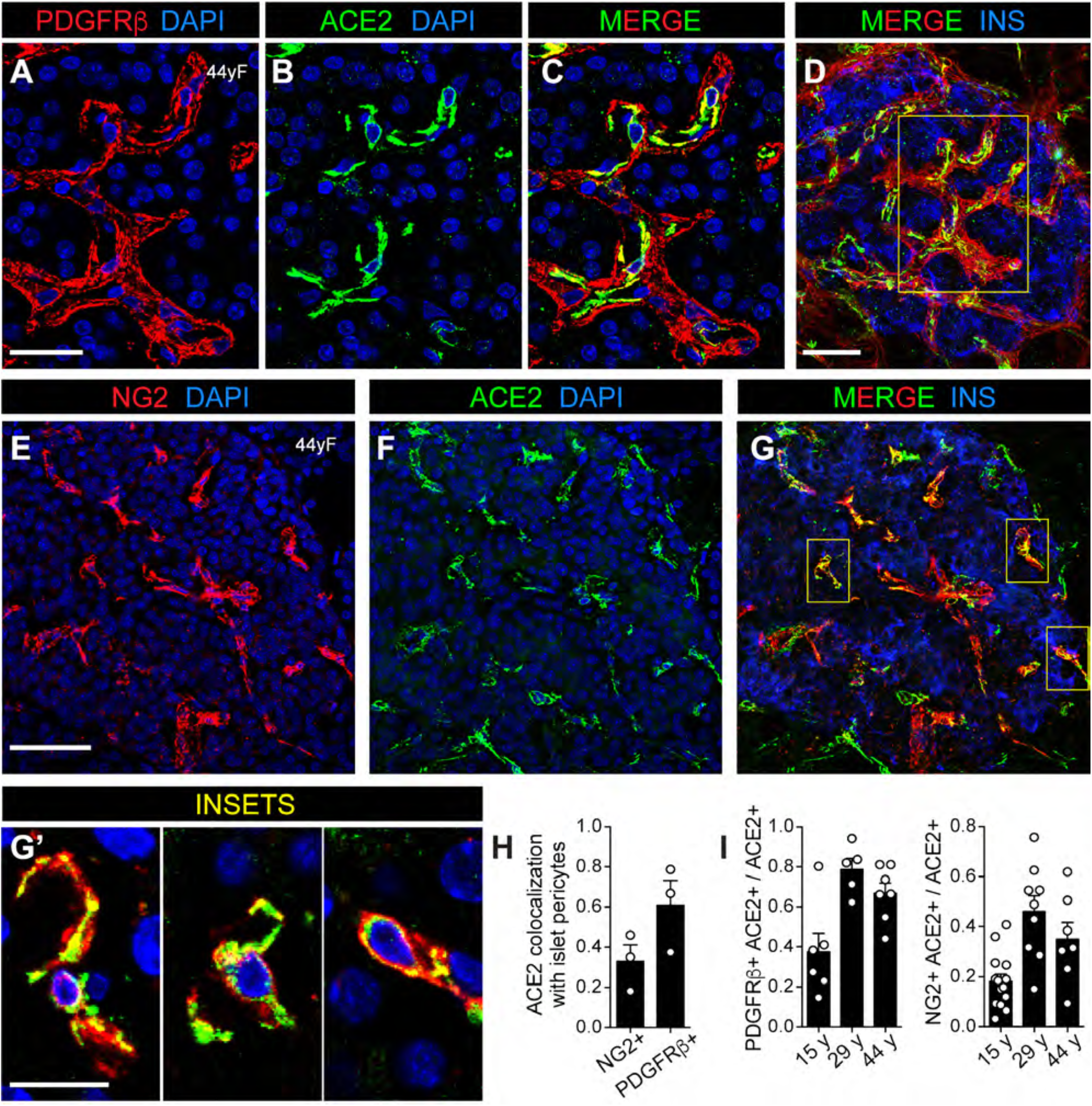
ACE2 Protein is Expressed by Islet Pericytes. (**A-C**) Maximum intensity projection of representative confocal images labelled for platelet-derived growth factor receptor β-positive cells (PDGFRβ, red) and ACE2 (green) in perivascular cells within a normal adult human islet; nuclear DAPI (blue). (**D**) Image of the same islet shown in A-C with insulin labeling (blue). Region within the yellow box in **D** is displayed in **A-C**. (**E-G**) Maximal intensity projection of pericyte labeling with an antibody against the pericyte marker neuron-glial antigen 2 (**E**, NG2, red) colocalized with ACE2 labeling (**F**, green) and nuclear DAPI (blue) in a normal adult human islet (**E-F**). (**G**) Image of the same islet shown in E-F with insulin labeling (blue). Regions within yellow boxes containing pericytes in **G** denote inset areas (**G’**) and represent a subset of islet ACE2-expressing pericytes. (**H**) Colocalization of ACE2 with NG2- or PDGFRβ-positive cells. (**I**) Quantification of ACE2-positive cells expressing pericyte markers NG2 and PDGFRβ Shown are average values obtained for 3 confocal planes per islet for a minimum of 5 islets per each donor of a given age. Scale bars are 20 μm (**A-C**), 40 μm (**D-G**), and 10 μm (**G’**). Human pancreatic donor information is available in Table S2 (**A-I,** donors HP1754, HP2041, HP2091). See also Figure S5.

### Both ACE2 and TMPRSS2 protein are present in pancreatic ducts but rarely are co-expressed

Our work indicated that TMPRSS2 protein was localized to the apical surface of intercalated and larger ducts in ND (**Figures 2B, 2C”, 2E, 2F”** and **S3E-H**), T2D (**Figures 2H, 2I”, 2K, 2L”** and **S3M-N**), and T1D pancreatic tissues (**Figures 2N, 2O”, 2Q, 2R”** and **S3S-V**). To further define ductal expression of TMPRSS2 and determine whether ACE2 was expressed beyond exocrine tissue capillaries, we visualized expression of these two proteins in relationship to cytokeratin-19 protein (KRT19), a ductal cell marker. TMPRSS2, but not ACE2, was expressed by KRT19-positive cells on the apical surface of intercalated and larger ducts throughout the exocrine compartment (**Figure 5A-D’**). Rarely, both ACE2 and TMPRSS2 were found on the apical surface of ductal epithelial cells but were spatially distinct (**Figure 5E-H’**). These findings agree with our scRNA-seq analysis in which *TMPRSS2* was more highly expressed in ductal cells than *ACE2*, but cells positive for both *ACE2* and *TMPRSS2* were rare. In addition, although *TMPRSS2* mRNA was detected in acinar cells by scRNA-seq (**Figure 1B**), TMPRSS2 protein was undetectable by immunofluorescence analysis (**Figure 5**), indicating that studies of mRNA and protein may be discordant.

**Figure 5.**
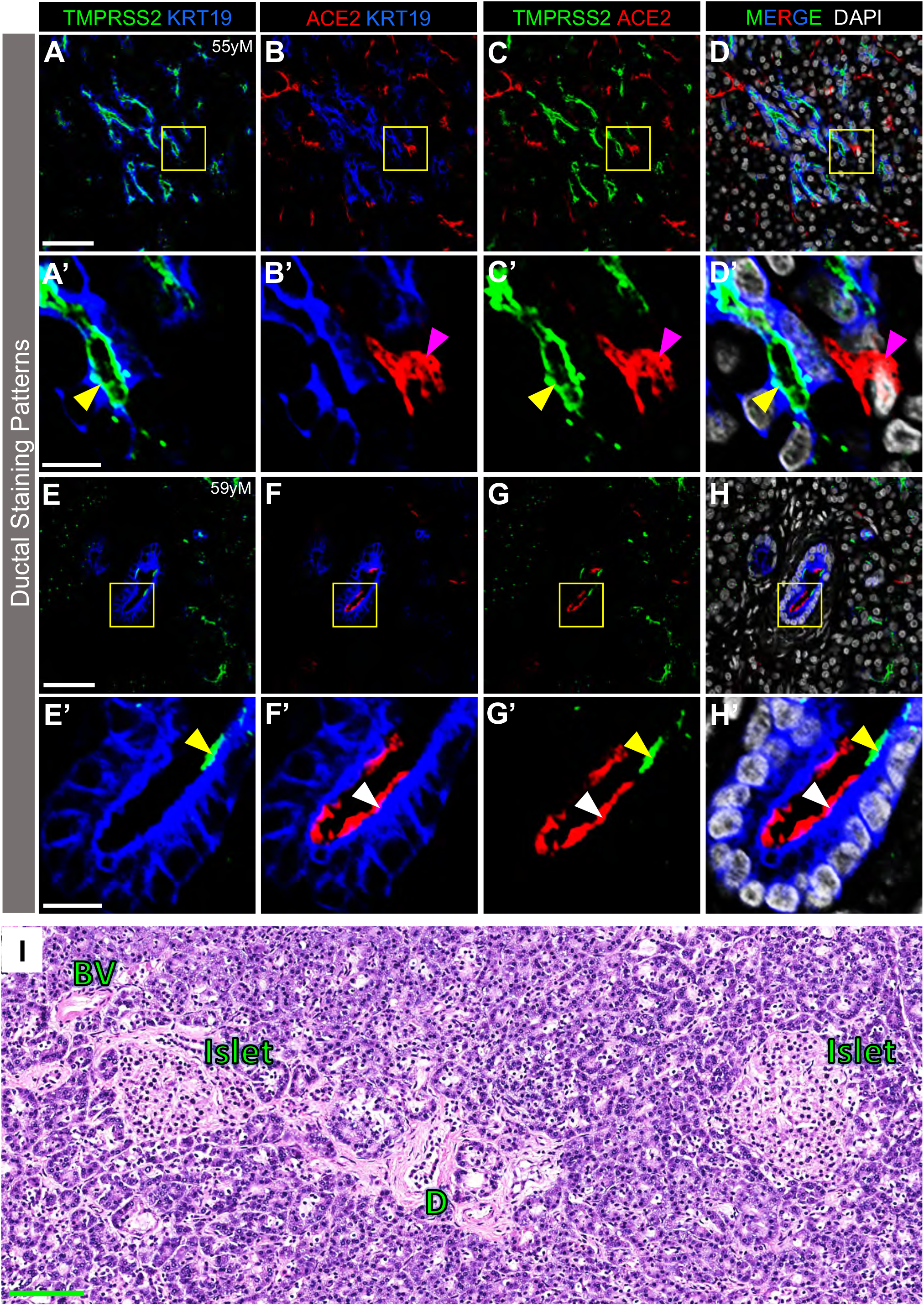
Analysis of ACE2 and TMPRSS2 Protein Expression in Pancreatic Ducts and Pancreatic Histology. (**A-D’**) TMPRSS2 (green), but not ACE2 (red; antibody R&D AF933), is expressed by KRT19-positive cells (blue) on the apical surface of intercalated and larger ducts throughout the exocrine compartment. DAPI (white). Yellow arrowheads point to TMPRSS2 on the apical surface of ductal cells (KRT19-positive). Magenta arrowheads point to ACE2-positive non-ductal cells. (**E-H’**) Rarely, both TMPRSS2 (green) and ACE2 (red; antibody R&D AF933) were expressed on the apical surface of ductal epithelial cells (KRT19, blue), but appeared to be spatially distinct. Yellow and white arrowheads point to spatially distinct TMPRSS2 and ACE2 labeling, respectively, localized to the apical surface of larger ducts (KRT19-positive cells). Inset areas (**A’-H’**) are marked by yellow boxes in **A-H**. Scale bars are 50 μm (**A-H**) and 10 μm (**A’-H’**). (**I**), Representative H&E image of human pancreas upon autopsy after COVID-19 disease. None of the seven donor samples evaluated showed signs of pancreatitis as determined by the histological absence of interstitial edema and inflammatory infiltrate, hemorrhage or necrosis. BV, blood vessel; D, duct. Scale bar in green is 100 μm. Human pancreatic donor information is available in Table S2 (**A-D’**, donor N8; **E-H’**, donor N9; **I**, COVID-19 donor 1). See also Figure S3.

Because rare cells in the exocrine compartment co-express ACE2 and TMPRSS2, it is possible that SARS-CoV-2 could infect those cells, induce pancreatitis, and consequently impact islet function. While recent case reports of patients with COVID-19 have noted mild elevations of amylase and lipase or frank pancreatitis (Karimzadeh et al., 2020; Meireles et al., 2020; Rabice et al., 2020; Wang et al., 2020), the vast majority of patients with COVID-19 have neither elevated pancreatic enzymes nor frank pancreatitis (Ashok et al., 2020; Bonney et al., 2020; Bruno et al., 2020). To investigate this possibility, we analyzed the histology of pancreatic tissue obtained at autopsy from seven COVID-19 positive patients, three of whom had diabetes. We did not find signs of pancreatitis, interstitial edema, inflammatory infiltrate, hemorrhage or necrosis (**Figure 5I**). It is important that exocrine inflammation and ACE2/TMPRSS2 expression be analyzed in a larger number of COVID-19 autopsy samples, particularly in those with new-onset diabetes. Such studies will be challenging since the pancreas rapidly undergoes autolysis upon death.

In summary, recent reviews, commentaries, and clinical guidelines (Apicella et al., 2020; Bornstein et al., 2020; Goldman et al., 2020; Gupta et al., 2020; Heaney et al., 2020) have highlighted an attractive hypothesis of direct cytotoxicity by SARS-CoV-2 to β cells. Here, we addressed the fundamental question of whether canonical SARS CoV-2 cell entry machinery is present in β cells of human pancreatic tissue. By examining pancreatic ACE2 and TMPRSS2 expression in normal donors and in those with diabetes, we found that, **1)** β cells do not coexpress *ACE2* and *TMPRSS2* transcripts by scRNA-seq analysis and neither protein is detectable in α or β cells by immunofluorescence; **2)** ACE2 is primarily expressed in the islet and exocrine tissue capillaries, including pericytes, as well as a subset of ductal cells; **3)** TMPRSS2 is primarily expressed in ductal cells; **4)** ACE2 and TMPRSS2 are infrequently co-expressed in pancreatic ducts; **5)** ACE2 and TMPRSS2 protein expression and distribution appear similar in ND and T2D pancreata but studies on additional pancreata are needed to definitively answer this question; and **6)** Alternative pathways facilitating SARS-CoV-2 viral infection in β cells cannot be excluded. These findings align considerably with independent observations from Kusmartseva and colleagues (Kusmartseva, 2020) and do not support the hypothesis that SARS-CoV-2 binds and infects islet β cells via a canonical pathway mediated by ACE2 and TMPRSS2, eliciting a diabetogenic effect. As outlined below in the “Limitations of the Study” section, additional studies are needed to investigate whether direct SARS-CoV-2 infection of β cells occurs or is detrimental to β cell health or function by other mechanisms. However, based on current data, it appears that the interaction of diabetes and SARS-CoV-2 is mediated by systemic inflammation and/or metabolic changes in other organs such as liver, muscle or adipose tissue.

### Limitations of the Study

We did not directly measure SARS-CoV-2 binding or entry into human β cells but instead assessed expression of canonical, co-receptors, ACE2 and TMPRSS2, in the human pancreas. While studies culturing SARS-CoV-2 and human islets *in vitro* will be of considerable interest, such experiments may be challenging to interpret as the islet isolation process itself may impact ACE2 or TMPRSS2 expression or susceptibility to viral infection. In addition, infection of islet cells within an intact, isolated islet with a variety of viruses has proven difficult as is selection of an appropriate ratio of viral particles and islet cells reflective of the *in vivo* environment. Furthermore, it is not clear if infection of an isolated islet accurately models the physiology of a vascularized, innervated islet within the context of the pancreas.

While the preponderance of research suggests that ACE2 and TMPRSS2 are the primary means for SARS-CoV-2 entry into host cells, knowledge about additional pathways and/or mechanisms of SARS-CoV-2 cell entry and infection is rapidly evolving. We sought, but did not find evidence, suggesting that other proposed pathways (namely *DPP4*, *CTSL*, *ADAM17*, *FURIN* or *TMPRSS4*) are involved in β cell entry.

We did not assess the presence of SARS-CoV-2, its proteins, or RNA in the pancreas from COVID-19-infected individuals. It is possible that SARS-CoV-2 could infect pancreatic exocrine cells, ductal cells, endothelial cells, or microvasculature, leading to pancreatic inflammation that negatively impacts β cell health or function. We did not find evidence of inflammation in the pancreas of seven COVID-19-infected patients (three of whom had diabetes). Future efforts are needed to collect a large number of pancreata from COVID-19-infected individuals with and without diabetes (long-standing and recent-onset) to search for presence of SARS-CoV-2 or inflammation (islet or exocrine). Since the pancreas undergoes rapid autolysis after death, care must be taken to collect the pancreas very soon after death for this analysis.

## Acknowledgements

We are especially thankful to organ donors and their families. This research was supported by funding provided by the National Institute of Diabetes and Digestive and Kidney Diseases, the Human Islet Research Network (HIRN; RRID:SCR_014393; https://hirnetwork.org; DK112232, DK123716, U01DK123594, DK104211, DK108120), and by DK106755, DK117147, DK117147-01A1S1, DK111757, DK112217, DK20593, and the Functional Genomics Core of DK19525. This manuscript used data acquired from and available at the Human Pancreas Analysis Program (HPAP-RRID:SCR_016202) Database (https://hpap.pmacs.upenn.edu), a HIRN consortium. This work was also supported by grants from the Doris Duke Charitable Foundation (DDCF 4043516256), Human Islet Research Network New Investigator Pilot Award (UC4 DK104162), JDRF, The Leona M. and Harry B. Helmsley Charitable Trust, and the Department of Veterans Affairs (BX000666). Human pancreatic islets were provided by the NIDDK-funded Integrated Islet Distribution Program at the City of Hope (NIH Grant # 2UC4 DK098085 RRID: SCR_014387; http://iidp.coh.org). Human kidney sections were provided by Dr. Agnes B. Fogo at Vanderbilt University Medical Center. We thank Amber M. Bradley for technical assistance.

## Author Contributions

K.C. J.C., M.B., and A.C.P. designed the experiments. K.C., J.C., M.B., and A.C.P. wrote the manuscript. K.C., J.C., S.S., W.W., L.G. J.A., M.E.K., M.F., A.M., C.D., D.S., R.B., R.J., R.W.S., K.L.H., G.V., M.B., and A.C.P. performed experiments or analyzed the data. All authors reviewed and edited the final manuscript. As the lead contact, A.C.P. is responsible for: 1) communication with the journal and co-authors; 2) addressing requests for reagents and resources; and 3) overseeing decisions and disputes related to the manuscript.

## Declaration of Interests

The authors declare no conflicts of interests.

## STAR METHODS

### Resource Availability

#### Lead contact

Further information and requests for resources and reagents should be directed to and will be fulfilled by the Lead Contact, Alvin C. Powers.

#### Materials Availability

This study did not generate new unique reagents.

#### Data and Code Availability

All data generated or analyzed during this study are included in this published article or in the data repositories listed in the Key Resources Table. Original/source data for Figure 1A (bulk RNA-seq of FACS sorted human islet α and β cells) is available under NCBI GEO accession numbers GSE67543 (Blodgett et al., 2015) and GSE57973 (Arda et al., 2016). Original/source data for Figure 1B and Figure S4J (single cell RNA-seq of human islets) is available under NCBI GEO accession number GSE84133 (Baron et al., 2016), GSE124742 (Camunas-Soler et al., 2020), and ArrayExpress (EBI) accession number E-MTAB-5061 (Segerstolpe et al., 2016). This manuscript used data acquired from the Human Pancreas Analysis Program (HPAP) Database (https://hpap.pmacs.upenn.edu), a Human Islet Research Network consortium.

### Experimental Model and Subject Details

#### Human Subjects

Pancreata and islets from juvenile, adult, T1D and T2D human donors were obtained through partnerships with the International Institute for Advancement of Medicine (IIAM), National Disease Research Interchange (NDRI), Integrated Islet Distribution Program (IIDP), and local organ procurement organizations. Pancreata from normal donors were processed either for islet isolation (Balamurugan et al., 2003) and/or histological analysis as described previously (described below and in Table S2) (Brissova et al., 2018; Dai et al., 2017). Pancreata from COVID-19 decedents after autopsy were obtained from the Translational Pathology Shared Resource (TPSR) at Vanderbilt University Medical Center (Nashville, TN) and and processed for histological analysis as according to standard procedures for clinical diagnostics (VUMC Histology Peloris Processing Protocol). Samples from donors of both sexes were used in our analyses. Donor demographic information is summarized in Table S2. The Vanderbilt University Institutional Review Board does not consider de-identified human pancreatic specimens to be human subjects research.

## METHOD DETAILS

### Human Pancreas Procurement and Preparation of Tissue for Histological Analysis

Pancreata from juvenile, adult, T1D and T2D donors were obtained within 18 hours from cross-clamp and maintained in cold preservation solution on ice until processing, as described previously (Balamurugan et al., 2003; Brissova et al., 2018). Pancreata recovered from seven COVID-19 decedants were obtained from Vanderbilt University Medical Center (Nashville, TN) within 8-27 hours of death. Donor demographic information is summarized in Table S2. Human kidney samples were provided by Dr. Agnes Fogo, Vanderbilt University. The Vanderbilt University Institutional Review Board has declared that studies on de-identified human pancreatic specimens do not quality as human subjects research.

### Mouse Tissue Preparation for Histological Analysis

Mice were maintained on standard rodent chow under a 12-hour light/dark cycle. Kidney from adult NOD.*Cg-Prkdc*^*scid*^*Il2rg*^*tm1Wjl*^/ Sz (NSG) mice ages 12 to 18 weeks (Jackson Laboratory) were isolated. Tissue specimens were processed for cryosections as described previously (Brissova et al., 2018).

### Immunohistochemical Analysis

Immunohistochemical analysis was performed on 5-μm cryosections (Figure 1C; Figure 3 and 4; Figure S2:A-I; S3A-D, I-L and O-R; S4 and S5) or formalin-fixed paraffin embedded (FFPE) pancreatic sections (Figure 2 and 5; Figure S2:J-W; S3E-H, M-N and S-V) as indicated and described previously (Brissova et al., 2014; Brissova et al., 2018; Wright et al., 2020). FFPE sections were first deparaffinized in xylene and ethanol followed by heat-induced epitope retrieval in a citrate-based antigen unmasking solution, pH 6.0 (see Key Resource Table). Immunofluorescence analysis of isolated islets was performed on 8-μm cryosections of islets embedded in collagen gels as described previously (Brissova et al., 2005; Brissova et al., 2018). Primary antibodies to all antigens and their working dilutions are listed in the Key Resource Table. The antigens were visualized using appropriate secondary antibodies listed in the Key Resource Table. Digital images were acquired with an Olympus FLUOVIEW FV3000 laser scanning confocal microscope (Olympus Corporation).

### Peptide Competition

We combined 1μg of anti-ACE2 antibody (ab15348) with or without 10μg of the immunizing human ACE2 peptide (ab15352) in antibody buffer solution (0.1% Triton X-100/1% BSA/1X PBS) and incubated overnight at 4°C with gentle agitation.

Immunofluorescence staining of 5-μm serial cryosections from the same donor was performed as described above. One section was incubated with the neutralized antibody buffer while the other section was incubated with the non-neutralized antibody buffer. ACE2 staining patterns were visualized via confocal microscopy as described above.

### H&E Staining

FFPE pancreas tissue sections (5-μm sections, n=7 donors) were processed according to standard procedures for clinical diagnostics (Vanderbilt University Medical Center Histology Peloris Processing Protocols). Slides were stained with hematoxylin & eosin and reviewed by a clinical pathologist (M.E.K.) at Vanderbilt University Medical Center.

### Bulk RNA-seq Data Acquisition

Transcripts Per Million (TPM) and Reads Per Kilobase Per Million (RPKM) normalized counts were extracted from publicly available RNA-seq datasets (Arda et al., 2016; Blodgett et al., 2015). The sources of the datasets are summarized in the Key Resources Table. GraphPad Prism v8 was used to generate plots in Figure 1A.

### Single Cell RNA-seq Data Acquisition

Raw gene count matrices were extracted from existing single cell RNA-seq datasets (Baron et al., 2016; Segerstolpe et al., 2016; Yang et al., 2020) and from the Human Pancreas Analysis Program (HPAP) Database (https://hpap.pmacs.upenn.edu), a Human Islet Research Network consortium. The sources of the datasets are summarized in the Key Resources Table. Gene count matrices were further analyzed using the R package Seurat version 3.1 (Stuart et al., 2019). Gene count measurements were normalized for each cell by library size and log-transformed using a size factor of 10,000 molecules per cell. The data was further scaled to unit variance and zero mean implemented in the “ScaleData” function. Cell types already annotated by the authors in the original study were used and thus, no clustering was performed. Seurat’s “DotPlot” function was used to generate plots shown in Figures 1B and S4J to visualize *ACE2* and *TMPRSS*2 scaled expression.

## QUANTIFICATION AND STATISTICAL ANALYSIS

### Statistical analysis

Bulk RNA-seq data are expressed as mean ± standard error of mean (Figures 1A). Sample size (n) is provided as the number of independent donor samples. A p-value less than 0.05 was considered significant. Statistical analysis (unpaired t-test) was performed using GraphPad Prism software. Mander’s coefficients were determined to quantify colocalization between pericyte markers (NG2 and PDGFRβ) and ACE2 in confocal images of islets using the ImageJ plugin “Just Another Co-localization Plugin”: (https://imagej.nih.gov/ij/plugins/track/jacop2.html).

## ADDITIONAL RESOURCES

None.

---

**Figure S1. Related to Figure 1.**
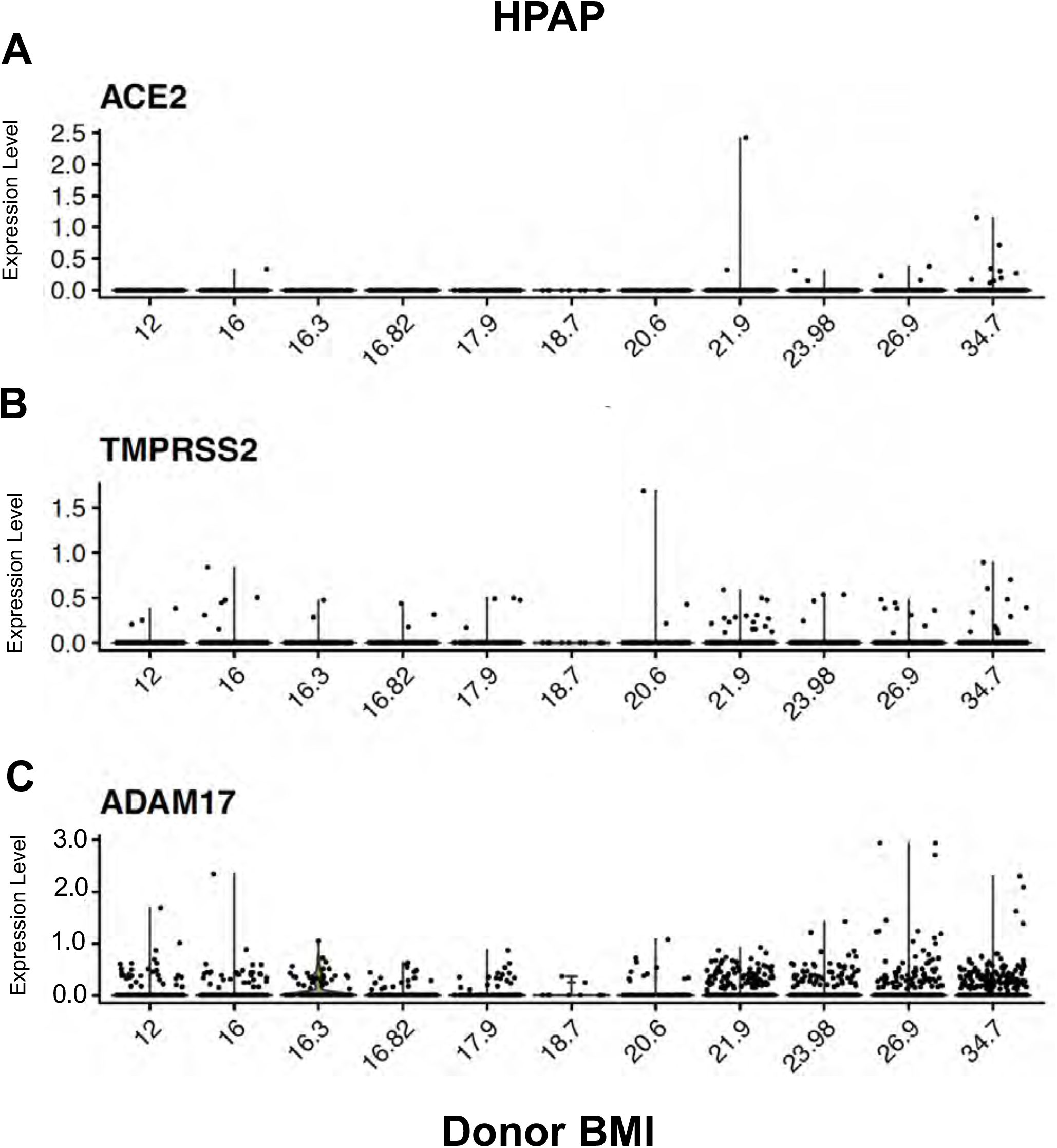
Stratification of *ACE2*, *TMPRSS2*, and *ADAM17* Expression in β cells by BMI. β cell gene expression from eleven donors (ages 1-39 years) from the HPAP scRNA-seq dataset (Kaestner et al., 2019) are displayed according to increasing BMI. Only one donor had an obese BMI. (**A-B**) *ACE2* and *TMPRSS2* expression in β cells did not show correlation with increasing BMI. (**C**) A trend towards increased *ADAM17* expression with BMI was identified. However, only one obese donor was available and analyzed in this dataset. Human pancreatic donor information is available in Table S2.

**Figure S2. Related to Figure 1 and 2.**
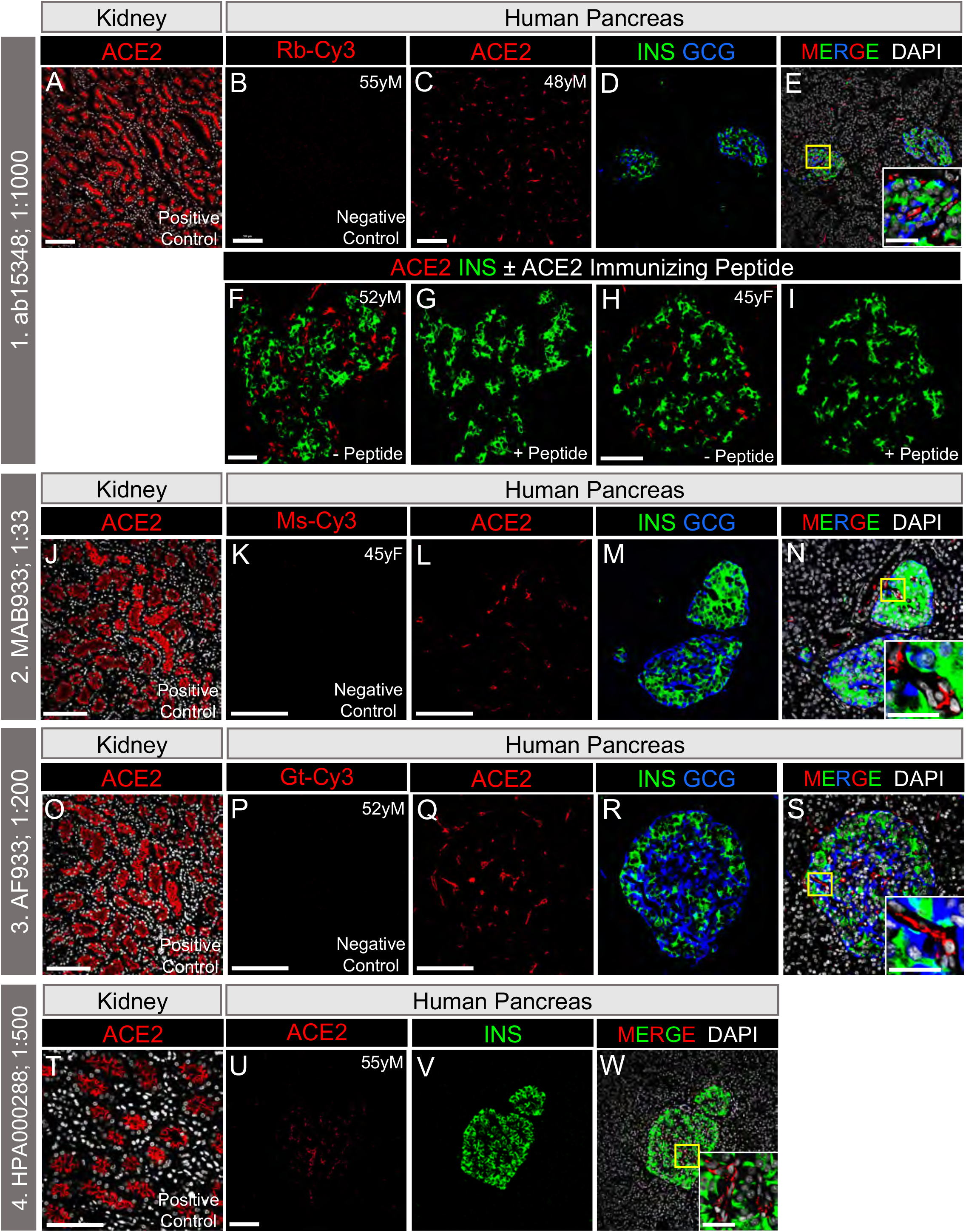
Testing and Characterization of Four ACE2-directed Antibodies on Human Pancreatic Tissue. (**A-E**) Characterization of ACE2 antibody (red, ab15348) used by Yang et al. (Yang et al., 2020). Antibody epitope encompasses the ACE2 C-terminal domain (human aa 788-805). Mouse kidney tissue served as a positive control for ACE2 (**A**), while normal adult human pancreatic tissue incubated with anti-rabbit-Cy3 secondary antibody only served as a negative control (**B**). Normal adult human pancreas labeled for ACE2 (red), INS (green, β cells) and GCG (blue, α cells) (**C-E**). Inset area is marked by a yellow box in MERGE column. (**F-I**) Competition with immunizing peptide neutralized ACE2 labeling, demonstrating the antibody’s antigen specificity. Scale bars are 100 μm (**A-E**) and 50 μm (Inset, **E** and **F-I**). (**J-N**) Characterization of ACE2 antibody (red, R&D MAB933) at same dilution (1:33) reported by Fignani et al. (preprint: Fignani, 2020). Antibody epitope encompasses the ACE2 extracellular domain (human aa 18-740). Human kidney tissue served as a positive control for ACE2 (**J**), while normal adult human pancreatic tissue incubated with anti-mouse-Cy3 secondary antibody only served as a negative control (**K**). Normal adult human pancreas labeled for ACE2 (red), INS (green, β cells) and GCG (blue, α cells) (**L-N**). Inset area is marked by a yellow box in MERGE column. Scale bars are 50 μm (**J-N**) and 25 μm (Inset, **N**). (**O-S**) Characterization of ACE2 antibody (red, R&D AF933) at same dilution (1:200) reported by Yang et al. (Yang et al., 2020). Antibody epitope encompasses the ACE2 extracellular domain (human aa 18-740). Human kidney served as a positive control for ACE2 (**O**), while normal adult human pancreatic tissue incubated with anti-goat-Cy3 secondary antibody only served as a negative control (**P**). Normal adult human pancreas labeled for ACE2 (red), INS (green, β cells) and GCG (blue, α cells) (**Q-S**). Inset area is marked by a yellow box in MERGE column. Scale bars are 50 μm (**O-R**) and 25 μm (Inset, **S**). (**T-W**) Characterization of ACE2 antibody (red, HPA000288) used by the Human Protein Atlas (Uhlen et al., 2015) and Hikmet et al. (Hikmet et al., 2020). Antibody epitope encompasses the ACE2 extracellular domain (human aa 1-111). Human kidney tissue served as a positive control for ACE2 (**T**). Normal adult human pancreas labeled for ACE2 (red) and INS (green, β cells) (**U-W**). Inset area is marked by a white dashed box in MERGE column. Scale bars are 100 μm (**T-V**) and 50 μm (Inset, **W**). DAPI (white). Human pancreatic donor information is available in Table S2 (**B**, donor N8; **C-E**, donor N4; **F-I**, donors N6 and N2; **J-N**, donor N2; **O-S**, donor N7; **T-W**, donor N8).

**Figure S3. Related to Figures 2 and 5.**
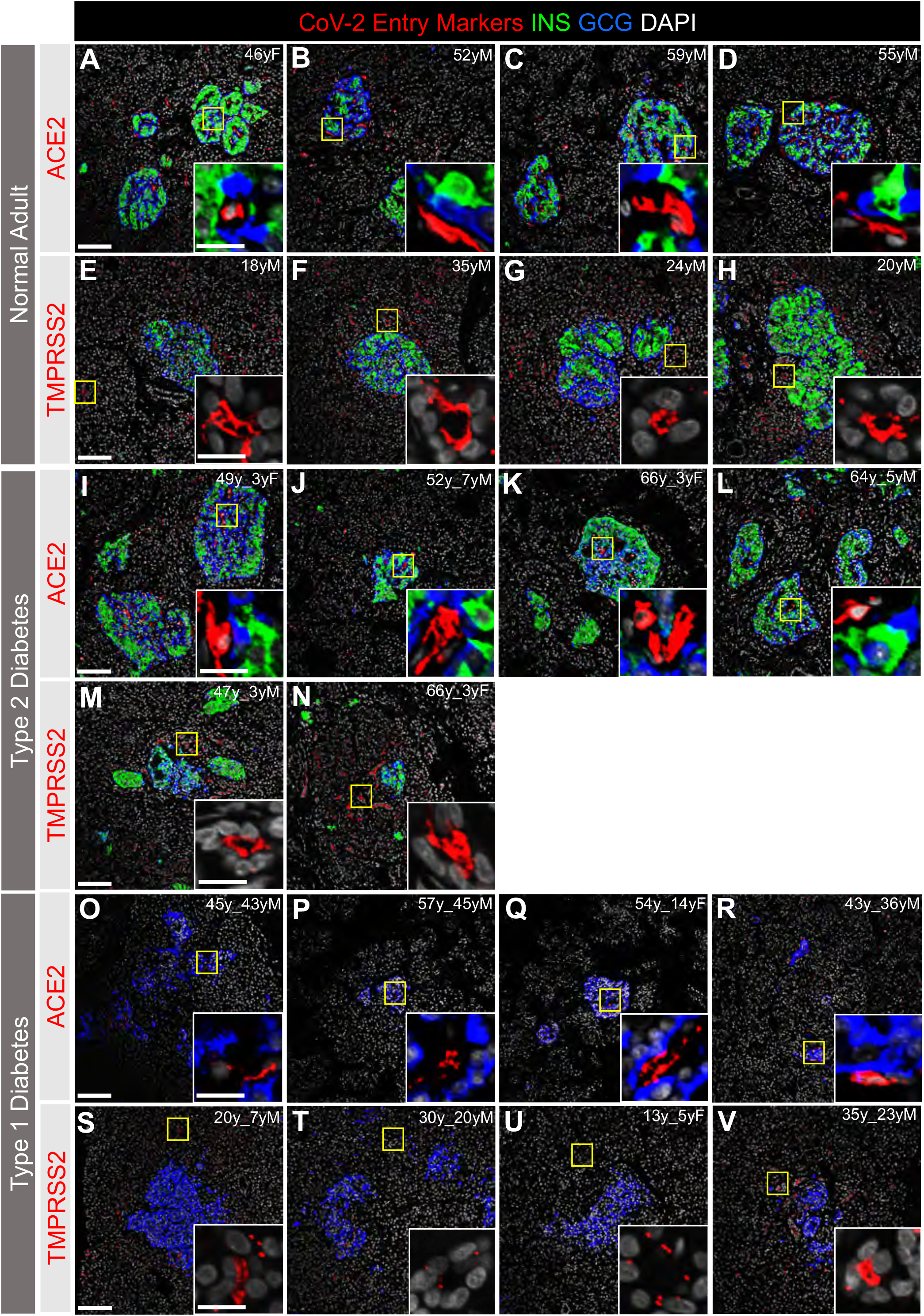
ACE2 and TMPRSS2 Protein are Not Detected by Immunofluorescence in α or β Cells from Normal, T2D or T1D Adult Donors. SARS-CoV-2 cell entry markers ACE2 (antibody ab15348) and TMPRSS2, both shown in red, are not detected in islet α cells (GCG, blue) or β cells (INS, green) in pancreatic sections from adult donors without diabetes (**A-H**) or donors with type 2 (**I-N**) or type 1 (**O-V**) diabetes. Insets are depicted by a yellow box. DAPI (white). Scale bars are 100 μm (**A-V**) and 25 μm (Insets). Human islet and pancreatic donor information is available in Table S2 (**A-D**, donors N3, N7, N9, N8; **E-H**, donors N14, N12, N11, N10; **I-L**, donors 2L, 2B, 2G, 2I; **M-N**, donors 2H, 2G; **O-R**, donors 1B, 1D, 1C, 1A; **S-V**, donors 1H, 1K, 1J, 1G).

**Figure S4. Related to Figures 1 and 2.**
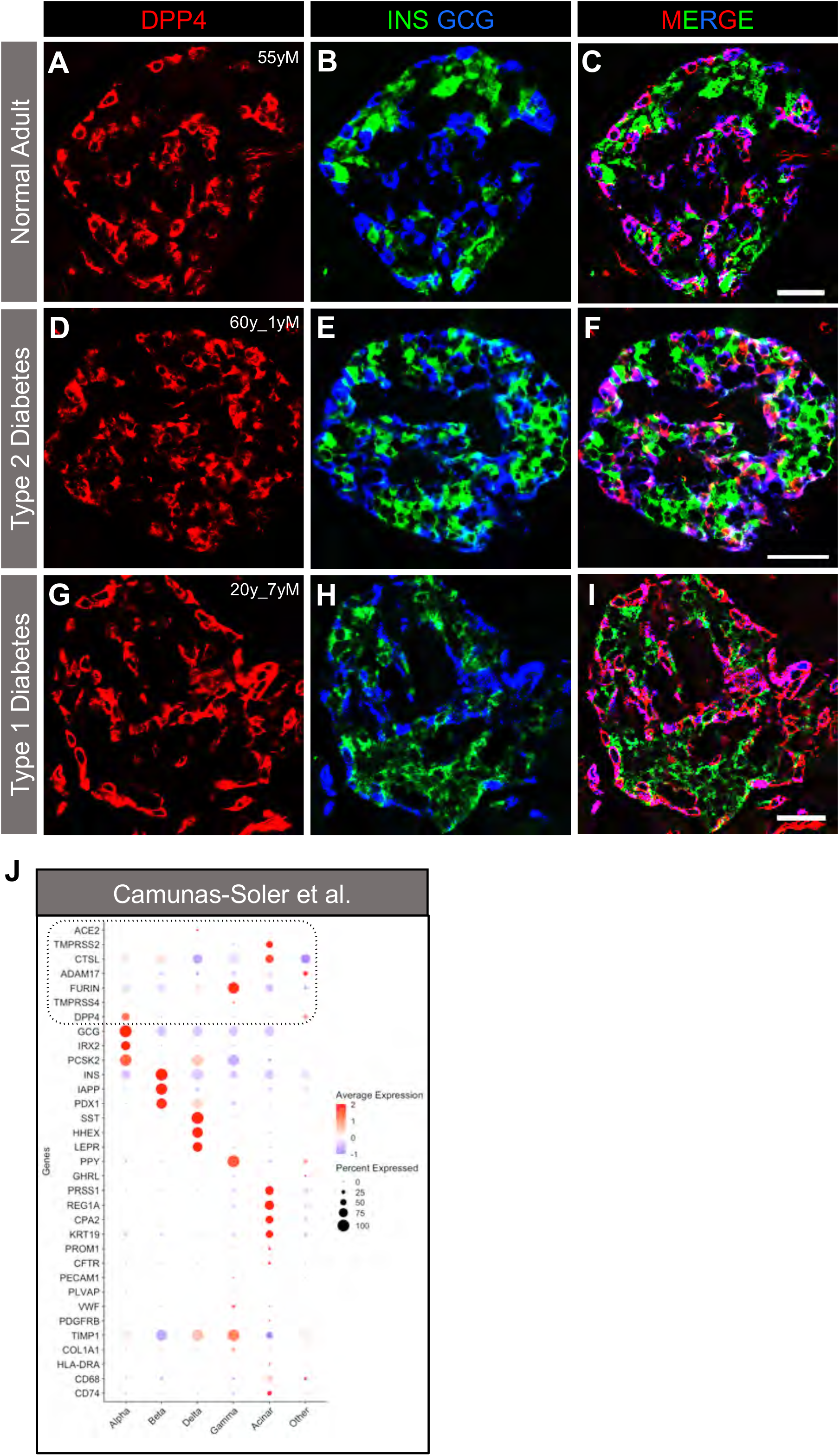
DPP4 is Expressed in α cells of Adult Normal and Diabetic Donors. DPP4 (red) is expressed in human pancreatic islet α cells (GCG, blue; Merged, magenta) but not β cells (INS, green) in pancreatic sections from adult donors without diabetes (**A-C**) or donors with type 2 (**D-F**) or type 1 (**G-I**) diabetes. Scale bars are 50 μm (**A-I**). Human islet and pancreatic donor information is available in Table S2 (**A-C**, donor N8; **D-F**, donor 2K; **G-I**, donor 1H). (**J**) Dot plots of *ACE2*, *TMPRSS2*, *CTSL*, *ADAM17*, *FURIN*, *TMPRSS4*, and *DPP4* expression compared with cell type-enriched genes from a previously published single cell (sc) RNA-seq datasets (Camunas-Soler et al., 2020). Dot size indicates percentage of cells in a given population expressing the gene; dot color represents scaled average expression. Dotted line highlights *ACE2*, *TMPRSS2*, *CTSL*, *ADAM17*, *FURIN*, *TMPRSS4*, and *DPP4* expression.

**Figure S5. Related to Figures 3 and 4.**
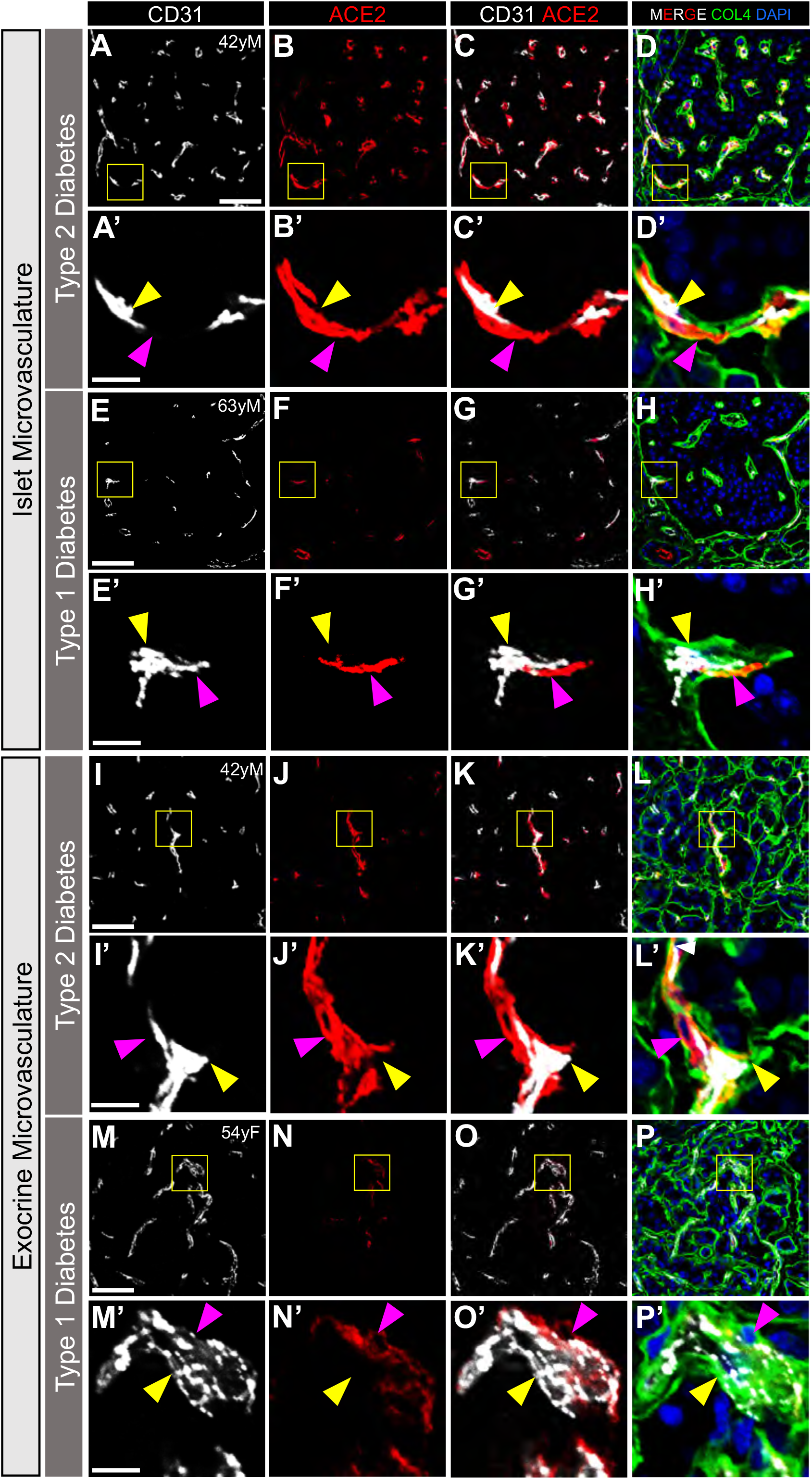
ACE2 is Localized to Islet and Exocrine Capillaries in Adult Human Pancreas of T2D and T1D donors. Representative images of endothelial cells (CD31, white) and ACE2-positive perivascular cells (red; antibody ab15348) in islet (**A-H’**) and exocrine tissue microvasculature (**I-P’**) of individuals with type 2 and type 1 diabetes. DAPI (white). Yellow arrowheads point to CD31-positive endothelial cells, while magenta arrowheads point to perivascular ACE2-positive cells. Insets (**A’-P’**) are depicted by yellow boxes in **A-P**. ACE2 positive perivascular cells abut extracellular matrix marker collagen-IV within the vascular basement membrane (collagen-IV [COL4], green; **D, D’, H, H’, L, L’, P, P’**). ACE2 labeling was reduced in both islets (**E-H’**) and exocrine tissue of individuals with type 1 diabetes (**M-P’**) compared to normal donors (Figure 3) and those with type 2 diabetes. Scale bars are 50 μm (**A-P**) and 10 μm (Insets, **A’-P’**). Human pancreatic donor information is available in Table S2 (**A-D’**, donor 2E; **E-H’**, donor 1F; **I-L’**, donor 2E; **M-P’**, donor 1C).

**Table S1. Related to Figure 1.** Number and Percentage of β cells that Express and Co-express Putative SARS-CoV-2 Cell Entry Genes Across Four Independent scRNA-seq Datasets.

| Genes | Droplet-based scRNA-seq |  |  |  | SMART-seq |  |  |  |
| --- | --- | --- | --- | --- | --- | --- | --- | --- |
| | HPAP<sup>a</sup><br>($\beta$ cell total, N=2828) | | Baron et al.<sup>b</sup><br>($\beta$ cell total, N=2525) | | Segerstolpe et al.<sup>c</sup><br>($\beta$ cell total, N=157) | | Camunas-Soler et al.<sup>c</sup><br>($\beta$ cell total, N=357) | |
| | # $\beta$ cell | % $\beta$ cells | # $\beta$ cell | % $\beta$ cells | # $\beta$ cell | % $\beta$ cells | # $\beta$ cell | % $\beta$ cells |
| *ACE2* | 17 | 0.6 | 4 | 0.2 | 3 | 1.9 | 12 | 3.4 |
| *TMPRSS2* | 60 | 2.1 | 7 | 0.3 | 4 | 2.5 | 2 | 0.6 |
| *TMPRSS4* | 0 | 0.0 | 0 | 0.0 | 0 | 0.0 | 9 | 2.5 |
| *CTSL* | 1421 | 50.2 | 977 | 38.7 | 132 | 84.1 | 291 | 81.5 |
| *FURIN* | 779 | 27.5 | 942 | 37.3 | 91 | 58.0 | 255 | 71.4 |
| *ADAM17* | 494 | 17.5 | 251 | 9.9 | 78 | 49.7 | 96 | 26.9 |
| *ACE2, TMPRSS2* | 0 | 0.0 | 0 | 0.0 | 0 | 0.0 | 0 | 0.0 |
| *ACE2, TMPRSS4* | 0 | 0.0 | 0 | 0.0 | 0 | 0.0 | 0 | 0.0 |
| *ACE2, CTSL* | 0 | 0.0 | 2 | 0.1 | 3 | 1.9 | 9 | 2.5 |
| *ACE2, FURIN* | 0 | 0.0 | 1 | 0.0 | 1 | 0.6 | 8 | 2.2 |
| *ACE2, ADAM17* | 0 | 0.0 | 0 | 0.0 | 1 | 0.6 | 3 | 0.8 |

<sup>a</sup>10x genomics; <sup>b</sup>InDrop (Klein et al., 2015); <sup>c</sup>SMART-seq2 (Picelli et al., 2014)

**Table S2.** Related to STAR Methods. Demographic Information of Donors.

| Donor ID |  | Age | Ethnicity / Race | Diabetes Duration | Sex | BMI | Cause of Death | Tissue/Islet Source |
| --- | --- | --- | --- | --- | --- | --- | --- | --- |
| **Juvenile (Histology)** | J1 | 5 days | Caucasian | -- | F | 14.9 | Anoxia | IIAM |
|  | J2 | 3 months | Caucasian | -- | M | 16.8 | Anoxia | NDRI |
|  | J3 | 10 months | Caucasian | -- | F | 15.4 | CVA | NDRI |
|  | J4 | 20 months | Caucasian | -- | F | 23.5 | Anoxia | IIAM |
|  | J5 | 5 years | Caucasian | -- | M | 16.2 | Anoxia | IIAM |
| **Normal Adult (Histology)** | N1 | 42 years | Caucasian | -- | M | 32.2 | Overdose | TNDS |
|  | N2 | 45 years | Caucasian | -- | F | 29.7 | Anoxia | OPO |
|  | N3 | 46 years | Caucasian | -- | F | 32.9 | CVA | IIAM |
|  | N4 | 48 years | Caucasian | -- | M | 24.6 | Anoxia | OPO |
|  | N5 | 51 years | Caucasian | -- | M | 20.4 | Anoxia | OPO |
|  | N6 | 52 years | Black | -- | M | 29.2 | ICH | TNDS |
|  | N7 | 52 years | Caucasian | -- | M | 28.1 | Head Trauma | OPO |
|  | N8 | 55 years | Black | -- | M | 35.6 | CVA | IIAM |
|  | N9 | 59 years | Caucasian | -- | M | 32.7 | Head Trauma | IIAM |
|  | N10 | 20 years | Hispanic | -- | M | 19.4 | Head Trauma | IIAM |
|  | N11 | 24 years | Caucasian | -- | M | 35.5 | ICH | IIAM |
|  | N12 | 35 years | Caucasian | -- | M | 26.8 | Head Trauma | IIAM |
|  | N13 | 20 years | Caucasian | -- | M | 27.8 | Head Trauma | NDRI |
|  | N14 | 18 years | Caucasian | -- | M | 25.1 | Head Trauma | IIAM |
|  | HP1754 | 15 years | N/A | -- | M | 22.6 | Head Trauma | IIAM |
|  | HP2041 | 29 years | N/A | -- | M | 22.3 | Head Trauma | IIAM |
|  | HP2091 | 44 years | N/A | -- | F | 23.7 | CVA | IIAM |
| **Adult T1D (Histology)** | 1A | 43 years | N/A | 36 years | M | 31.2 | CVA | NDRI |
|  | 1B | 45 years | Caucasian | 43 years | M | 25.0 | Anoxia | IIAM |
|  | 1C | 54 years | Caucasian | 14 years | F | 24.9 | Anoxia | IIAM |
|  | 1D | 57 years | Black | 45 years | M | 33.3 | CVA | IIAM |
|  | 1E | 58 years | Caucasian | 31 years | M | 21.8 | Anoxia | NDRI |
|  | 1F | 63 years | Caucasian | 44 years | M | 24.1 | Anoxia | IIAM |
|  | 1G | 35 years | Caucasian | 23 years | M | 26.9 | Anoxia | NDRI |
|  | 1H | 20 years | Caucasian | 7 years | M | 25.5 | Anoxia | NDRI |
|  | 1I | 27 years | Caucasian | 17 years | M | 18.4 | Anoxia | NDRI |
|  | 1J | 13 years | Caucasian | 5 years | M | 19.1 | Anoxia | IIAM |
|  | 1K | 30 years | Caucasian | 20 years | M | 29.8 | Anoxia | NDRI |
| **Adult T2D (Histology)** | 2A | 44 years | Caucasian | 7 years | M | 44.4 | CVA | IIAM |
|  | 2B | 52 years | Caucasian | 7 years | M | 33.6 | CVA | IIAM |
|  | 2C | 52 years | Asian | 10 years | F | 21.9 | CVA | NDRI |
|  | 2D | 52 years | Caucasian | < 1 year | F | 29.2 | CVA | IIAM |
|  | 2E | 42 years | Black | < 1 year | M | 42.0 | CVA | IIAM |
|  | 2F | 43 years | Black | 1 year | M | 36.0 | Head Trauma | IIAM |
|  | 2G | 66 years | Caucasian | 3 years | F | 32.8 | CVA | IIAM |
|  | 2H | 47 years | Caucasian | 3 years | M | 31.3 | CVA | IIAM |
|  | 2I | 64 years | Caucasian | 5 years | M | 33.2 | ICH | IIAM |
|  | 2J | 59 years | Caucasian | 6 years | F | 27.5 | CVA | IIAM |
|  | 2K | 60 years | Caucasian | 1 year | M | 38.3 | CVA | IIAM |
|  | 2L | 49 years | Caucasian | 3 years | F | 33.8 | CVA | IIAM |
| **Normal Adult Islets (Gels and scRNA-Seq)** | I1 | 40 years | Caucasian | -- | F | 30.8 | Head Trauma | IIDP |
|  | I2 | 41 years | N/A | -- | M | 20.3 | N/A | IIDP |
|  | I3 | 42 years | Caucasian | -- | M | 32.2 | Overdose | IIDP |
|  | HPAP022 | 39 years | Caucasian | -- | F | 34.7 | Anoxia | HPAP |
|  | HPAP026 | 24 years | Caucasian | -- | M | 20.8 | Anoxia | HPAP |
|  | HPAP034 | 13 years | Caucasian | -- | M | 18.6 | Head Trauma | HPAP |
|  | HPAP035 | 35 years | Caucasian | -- | M | 26.9 | Anoxia | HPAP |
|  | HPAP036 | 23 years | Caucasian | -- | F | 16 | Head Trauma | HPAP |
|  | HPAP037 | 35 years | Caucasian | -- | F | 21.9 | CVA | HPAP |
|  | HPAP039 | 5 years | Caucasian | -- | F | 16.3 | Anoxia | HPAP |
|  | HPAP040 | 35 years | Caucasian | -- | M | 23.9 | CVA | HPAP |
|  | HPAP042 | 1 year | Caucasian | -- | M | 17.9 | Anoxia | HPAP |
|  | HPAP044 | 3 years | Caucasian | -- | F | 12 | Anoxia | HPAP |
|  | HPAP047 | 8 years | Caucasian | -- | M | 16.8 | CVA | HPAP |
| **COVID-19 Patient Autopsy Samples (Histology)** | 1 | 82 years | Caucasian | -- | M | 26.8 | ALI | VUMC Autopsy |
|  | 2 | 97 years | Caucasian | -- | F | 19.7 | ALI | VUMC Autopsy |
|  | 3 | 81 years | Caucasian | >10 years<sup>a</sup> | M | 23.3 | ALI | VUMC Autopsy |
|  | 4 | 60 years | Hispanic | -- | M | 36.7 | ALI | VUMC Autopsy |
|  | 5 | 51 years | Hispanic | 23 years | M | 29.4 | ALI | VUMC Autopsy |
|  | 6 | 60 years | Caucasian | -- | F | 38.4 | PE | VUMC Autopsy |
|  | 7 | 71 years | Black | Pre-existing<sup>b</sup> | M | 31.5 | ALI | VUMC Autopsy |

ALI – acute lung injury; CVA, cerebrovascular accident; HPAP – Human Pancreas Analysis Program (Human Islet Research Network); ICH, intracerebral hemorrhage; IIAM – International Institute for the Advancement of Medicine; IIDP – Integrated Islet Distribution Program; N/A – not available; NDRI – National Disease Research Interchange; OPO – Organ Procurement Organization; PE – pulmonary embolism; T1D = type 1 diabetes; T2D – type 2 diabetes; TNDS – Tennessee Donor Services, Nashville; VUMC Autopsy – Vanderbilt University Medical Center Autopsy Pathology
<sup>a</sup>Oldest clinical patient note including diagnosis of diabetes mellitus was signed in 2010, suggesting disease duration of at least 10 years.
<sup>b</sup>Patient was prescribed an oral anti-diabetic medication confirming pre-existing diabetes diagnosis of unknown duration prior to admission with COVID-19.

## KEY RESOURCES TABLE

| REAGENT or RESOURCE | SOURCE | IDENTIFIER |
| --- | --- | --- |
| **Antibodies** |  |  |
| Mouse anti-Cytokeratin 19 (1:1000) | Abcam | Cat# ab194399 |
| Rabbit anti-TMPRSS2 (1:400) | Sigma | Cat# HPA035787, RRID:AB_2674782 |
| Mouse anti-Glucagon (1:500-1:1000) | Abcam | Cat# ab10988, RRID:AB_297642 |
| Guinea pig anti-Glucagon (1:500) | LSBio | Cat# LS-C202759-50 |
| Rat anti-C-peptide (1:1000) | DSHB | Cat# GN-ID4, RRID:AB_2255626 |
| Guinea pig anti-Insulin (1:500-1:1000) | Dako | Cat# A0564, RRID:AB_2617169 |
| Rabbit anti-ACE2 (1:500) | Atlas Antibodies | Cat# HPA000288, RRID:AB_1078160 |
| Rabbit anti-ACE2 (1:1000) | Abcam | Cat# ab15348, RRID:AB_301861 |
| Goat anti-ACE2 (1:200, Figure S1,3A-E; 1:400, Figure 4) | R&D | Cat# AF933, RRID:AB_355722 |
| Mouse anti-ACE2 (1:33) | R&D | Cat# MAB933, RRID:AB_2223153 |
| Mouse anti-CD31 (1:50) | BD Biosciences | Cat# 550389, RRID:AB_2252087 |
| Goat anti-collagen IV alpha 1 (1:500) | Novus | Cat# NBP1-26549, RRID:AB_1853202 |
| Goat anti-DPP4 (1:200) | R&D | Cat# AF1180, RRID:AB_354651 |
| Mouse anti-NG2 [647] (1:50) | R&D | Cat# FAB2585R; |
| Goat anti-PDGFRβ (1:100) | R&D | Cat# AF385, RRID:AB_355339 |
| Donkey anti-mouse-Cy2 (1:500) | Jackson ImmunoResearch | Cat# 715-225-150, RRID:AB_2340826 |
| Donkey anti-mouse-Cy3 (1:500) | Jackson ImmunoResearch | Cat# 715-165-150, RRID:AB_2340813 |
| Donkey anti-mouse-Cy5 (1:250 – 1:500) | Jackson ImmunoResearch | Cat# 715-175-151, RRID:AB_2340820 |
| Donkey anti-rabbit-Cy3 (1:500) | Jackson ImmunoResearch | Cat# 711-165-152, RRID:AB_2307443 |
| Donkey anti-rat-Cy2 (1:500) | Jackson ImmunoResearch | Cat# 712-225-153, RRID:AB_2340674 |
| Donkey anti-guinea pig-Cy2 (1:500) | Jackson ImmunoResearch | Cat# 706-225-148, RRID:AB_2340467 |
| Donkey anti-goat-Cy2 (1:500) | Jackson ImmunoResearch | Cat# 705-225-147, RRID:AB_2307341 |
| Donkey anti-goat-Cy3 (1:500) | Jackson ImmunoResearch | Cat# 705-165-147, RRID:AB_2307351 |
| Donkey anti-guinea pig-Cy5 (1:250-1:500) | Jackson ImmunoResearch | Cat# 706-175-148, RRID:AB_2340462 |
| Normal Donkey Serum | Jackson ImmunoResearch | Cat# 017-000-121, RRID: AB_2337258 |
| **Bacterial and Virus Strains** |  |  |
| n/a |  |  |
| **Biological Samples** |  |  |
| Human pancreatic islets | Integrated Islet Distribution Program (IIDP) | http://iidp.coh.org,<br>RRID: SCR_014387 |
| Human pancreatic tissue | International Institute for the Advancement of Medicine (IIAM) | http://www.iiam.org,<br>RRID: SCR_016172 |
| Human pancreatic tissue | National Disease Research Interchange (NDRI) | http://ndriresource.org,<br>RRID: SCR_000550 |
| **Chemicals, Peptides, and Recombinant Proteins** |  |  |
| Propidium Iodide | BD Biosciences | Cat# 556463 |
| Antigen Unmasking Solution, Citrate-Based | Vector Laboratories | Cat# H-3300-250 |
| 1X PBS | Corning | Cat# 46-013-CM |
| Triton X-100 | USB | Cat# 22686 |
| BSA | Sigma | Cat# A6003-25G |
| Human ACE2 Peptide | Abcam | Cat# ab15352 |
| **Critical Commercial Assays** |  |  |
| RNAqueous-Micro Kit | Fisher Scientific (Invitrogen) | Cat# AM1931 |
| High Capacity cDNA Reverse Transcription Kit | Fisher Scientific (Invitrogen) | Cat# 4368814 |
| **Deposited Data** |  |  |
| RNA-seq data for FACS purified human alpha and beta cells (Blodgett et al., 2015) | NCBI Gene Expression Omnibus | GEO: GSE67543 |
| RNA-seq data for FACS purified human alpha and beta cells (Arda et al., 2016) | NCBI Gene Expression Omnibus | GEO: GSE57973 |
| Single-cell RNA-seq data for human islets (Baron et al., 2016) | NCBI Gene Expression Omnibus | GEO: GSE84133 |
| Single-cell RNA-seq data for human islets (Camunas-Soler et al., 2020) | NCBI Gene Expression Omnibus | GEO: GSE124742 |
| Single-cell RNA-seq data for human islets (Segerstolpe et al., 2016) | ArrayExpress | E-MTAB-5061 |
| Single-cell RNA-seq data for human islets (Human Pancreas Analysis Program [HPAP]) | HPAP Database | https://hpap.pmacs.upenn.edu<br>RRID:SCR_014393 |
| **Experimental Models: Cell Lines** |  |  |
| n/a |  |  |
| **Experimental Models: Organisms/Strains** |  |  |
| n/a |  |  |
| **Recombinant DNA** |  |  |
| n/a |  |  |
| **Software and Algorithms** |  |  |
| FV31S-SW Viewer 2.4.1.198 | Olympus | https://www.olympus-lifescience.com/en/support/downloads/ |
| Seurat 3.1 R package | (Butler et al., 2018)<br>(Stuart et al., 2019) | N/A |
| Prism for macOS v.8.4.3 | Graphpad Software | http://www.graphpad.com,<br>RRID: SCR_002798 |
| JACoP: Just Another Co-localization Plugin | ImageJ | https://imagej.nih.gov/ij/plugins/track/jacop2.html |

## Notes

### Competing Interest Statement

The authors have declared no competing interest.

### Summary of Updates

We have included new and revised experimental data as follows: Figure 1B, Figure 2, Figure 4, Figure 5I, Figure S1, Figure S2F-I, Figure S4J and Table S1.

## REFERENCES

Aid, M., Busman-Sahay, K., Vidal, S., Maliga, Z., Bondoc, S., Starke, C., Terry, M., Jacobson, C., Wrijil, L., Ducat, S., Brook, O., Miller, A., Porto, M., Pellegrini, K., Pino, M., Hoang, T., Chandrashekar, A., Patel, S., Stephenson, K., Bosinger, S., Andersen, H., Lewis, M., Hecht, J., Sorger, P., Martinot, A., Estes, J., Barouch, D. (2020). Vascular disease and thrombosis in SARS-CoV-2 infected Rhesus macaques. Cell *In Press*.

Almaca, J., Caicedo, A., and Landsman, L. (2020). Beta cell dysfunction in diabetes: the islet microenvironment as an unusual suspect. Diabetologia 63, 2076–2085.

Almaca, J., Weitz, J., Rodriguez-Diaz, R., Pereira, E., and Caicedo, A. (2018). The pericyte of the pancreatic islet regulates capillary diameter and local blood flow. Cell Metab 27, 630–644 e634.

Apicella, M., Campopiano, M.C., Mantuano, M., Mazoni, L., Coppelli, A., and Del Prato, S. (2020). COVID-19 in people with diabetes: understanding the reasons for worse outcomes. Lancet Diabetes Endocrinol.

Arda, H.E., Li, L., Tsai, J., Torre, E.A., Rosli, Y., Peiris, H., Spitale, R.C., Dai, C., Gu, X., Qu, K., et al. (2016). Age-dependent pancreatic gene regulation reveals mechanisms governing human beta cell function. Cell Metab 23, 909–920.

Ashok, A., Faghih, M., and Singh, V.K. (2020). Mild pancreatic enzyme elevations in COVID-19 pneumonia: synonymous with injury or noise? Gastroenterology.

Balamurugan, A.N., Gu, Y., Miyamoto, M., Wang, W., Inoue, K., and Tabata, Y. (2003). Isolation, culture, and functional characteristics of diabetic islets [corrected]. Pancreas 26, 102–103.

Baron, M., Veres, A., Wolock, S.L., Faust, A.L., Gaujoux, R., Vetere, A., Ryu, J.H., Wagner, B.K., Shen-Orr, S.S., Klein, A.M., et al. (2016). A single-cell transcriptomic map of the human and mouse pancreas reveals inter- and intra-cell population structure. Cell Syst 3, 346–360 e344.

Barron, E., Bakhai, C., Kar, P., Weaver, A., Bradley, D., Ismail, H., Knighton, P., Holman, N., Khunti, K., Sattar, N., et al. (2020). Associations of type 1 and type 2 diabetes with COVID-19-related mortality in England: a whole-population study. Lancet Diabetes Endocrinol 8, 813–822.

Blodgett, D.M., Nowosielska, A., Afik, S., Pechhold, S., Cura, A.J., Kennedy, N.J., Kim, S., Kucukural, A., Davis, R.J., Kent, S.C., et al. (2015). Novel observations from next-generation RNA sequencing of highly purified human adult and fetal islet cell subsets. Diabetes 64, 3172–3181.

Bonney, G.K., Gao, Y., Chew, C.A., and Windsor, J.A. (2020). SARS-COV-2 associated acute pancreatitis: Cause, consequence or epiphenomenon? Pancreatology.

Bornstein, S.R., Rubino, F., Khunti, K., Mingrone, G., Hopkins, D., Birkenfeld, A.L., Boehm, B., Amiel, S., Holt, R.I., Skyler, J.S., et al. (2020). Practical recommendations for the management of diabetes in patients with COVID-19. Lancet Diabetes Endocrinol 8, 546–550.

Breidenbach, J.D., Dube, P., Ghosh, S., Abdullah, B.N., Modyanov, N.N., Malhotra, D., Dworkin, L.D., Haller, S.T., and Kennedy, D.J. (2020). Impact of Comorbidities on SARS-CoV-2 Viral Entry-Related Genes. J Pers Med 10.

Brissova, M., Aamodt, K., Brahmachary, P., Prasad, N., Hong, J.Y., Dai, C., Mellati, M., Shostak, A., Poffenberger, G., Aramandla, R., et al. (2014). Islet microenvironment, modulated by vascular endothelial growth factor-A signaling, promotes beta cell regeneration. Cell Metab 19, 498–511.

Brissova, M., Fowler, M.J., Nicholson, W.E., Chu, A., Hirshberg, B., Harlan, D.M., and Powers, A.C. (2005). Assessment of human pancreatic islet architecture and composition by laser scanning confocal microscopy. J Histochem Cytochem 53, 1087–1097.

Brissova, M., Haliyur, R., Saunders, D., Shrestha, S., Dai, C., Blodgett, D.M., Bottino, R., Campbell-Thompson, M., Aramandla, R., Poffenberger, G., et al. (2018). alpha Cell Function and Gene Expression Are Compromised in Type 1 Diabetes. Cell Rep 22, 2667–2676.

Bruno, G., Fabrizio, C., Santoro, C.R., and Buccoliero, G.B. (2020). Pancreatic injury in the course of coronavirus disease 2019: A not-so-rare occurrence. J Med Virol.

Camunas-Soler, J., Dai, X.Q., Hang, Y., Bautista, A., Lyon, J., Suzuki, K., Kim, S.K., Quake, S.R., and MacDonald, P.E. (2020). Patch-seq links single-cell transcriptomes to human islet dysfunction in diabetes. Cell Metab 31, 1017–1031 e1014.

Cariou, B., Hadjadj, S., Wargny, M., Pichelin, M., Al-Salameh, A., Allix, I., Amadou, C., Arnault, G., Baudoux, F., Bauduceau, B., et al. (2020). Phenotypic characteristics and prognosis of inpatients with COVID-19 and diabetes: the CORONADO study. Diabetologia 63, 1500–1515.

Chee, Y.J., Ng, S.J.H., and Yeoh, E. (2020). Diabetic ketoacidosis precipitated by Covid-19 in a patient with newly diagnosed diabetes mellitus. Diabetes Res Clin Pract 164, 108166.

Chen, L., Li, X., Chen, M., Feng, Y., and Xiong, C. (2020). The ACE2 expression in human heart indicates new potential mechanism of heart injury among patients infected with SARS-CoV-2. Cardiovasc Res 116, 1097–1100.

Clausen, T.M., Sandoval, D.R., Spliid, C.B., Pihl, J., Perrett, H.R., Painter, C.D., Narayanan, A., Majowicz, S.A., Kwong, E.M., McVicar, R.N., et al. (2020). SARS-CoV-2 Infection Depends on Cellular Heparan Sulfate and ACE2. Cell.

Dai, C., Hang, Y., Shostak, A., Poffenberger, G., Hart, N., Prasad, N., Phillips, N., Levy, S.E., Greiner, D.L., Shultz, L.D., et al. (2017). Age-dependent human beta cell proliferation induced by glucagon-like peptide 1 and calcineurin signaling. J Clin Invest 127, 3835–3844.

Drucker, D.J. (2020). Coronavirus infections and type 2 diabetes-shared pathways with therapeutic implications. Endocr Rev 41.

Filippi, C.M., and von Herrath, M.G. (2008). Viral trigger for type 1 diabetes: pros and cons. Diabetes 57, 2863–2871.

Goldman, N., Fink, D., Cai, J., Lee, Y.N., and Davies, Z. (2020). High prevalence of COVID-19-associated diabetic ketoacidosis in UK secondary care. Diabetes Res Clin Pract 166, 108291.

Gupta, A., Madhavan, M.V., Sehgal, K., Nair, N., Mahajan, S., Sehrawat, T.S., Bikdeli, B., Ahluwalia, N., Ausiello, J.C., Wan, E.Y., et al. (2020). Extrapulmonary manifestations of COVID-19. Nat Med 26, 1017–1032.

Hamming, I., Timens, W., Bulthuis, M.L., Lely, A.T., Navis, G., and van Goor, H. (2004). Tissue distribution of ACE2 protein, the functional receptor for SARS coronavirus. A first step in understanding SARS pathogenesis. J Pathol 203, 631–637.

He, L., Mae, M., Sun, Y., Muhl, L, Nahar, K., et al (2020). Pericyte-specific vascular expression of SARS-CoV-2 receptor ACE2 – implications for microvascular inflammation and hypercoagulopathy in COVID-19 patients. Biorxiv.

Heaney, A.I., Griffin, G.D., and Simon, E.L. (2020). Newly diagnosed diabetes and diabetic ketoacidosis precipitated by COVID-19 infection. Am J Emerg Med.

Hikmet, F., Mear, L., Edvinsson, A., Micke, P., Uhlen, M., and Lindskog, C. (2020). The protein expression profile of ACE2 in human tissues. Mol Syst Biol 16, e9610.

Hoffmann, M., Kleine-Weber, H., Schroeder, S., Kruger, N., Herrler, T., Erichsen, S., Schiergens, T.S., Herrler, G., Wu, N.H., Nitsche, A., et al. (2020). SARS-CoV-2 cell entry depends on ACE2 and TMPRSS2 and is blocked by a clinically proven protease inhibitor. Cell 181, 271–280 e278.

Hollstein, T., Schulte, D.M., Schulz, J., Gluck, A., Ziegler, A.G., Bonifacio, E., Wendorff, M., Franke, A., Schreiber, S., Bornstein, S.R., et al. (2020). Autoantibody-negative insulin-dependent diabetes mellitus after SARS-CoV-2 infection: a case report. Nat Metab.

Holman, N., Knighton, P., Kar, P., O’Keefe, J., Curley, M., Weaver, A., Barron, E., Bakhai, C., Khunti, K., Wareham, N.J., et al. (2020). Risk factors for COVID-19-related mortality in people with type 1 and type 2 diabetes in England: a population-based cohort study. Lancet Diabetes Endocrinol 8, 823–833.

Kaestner, K.H., Powers, A.C., Naji, A., Consortium, H., and Atkinson, M.A. (2019). NIH Initiative to Improve Understanding of the Pancreas, Islet, and Autoimmunity in Type 1 Diabetes: The Human Pancreas Analysis Program (HPAP). Diabetes 68, 1394–1402.

Karimzadeh, S., Manzuri, A., Ebrahimi, M., and Huy, N.T. (2020). COVID-19 presenting as acute pancreatitis: Lessons from a patient in Iran. Pancreatology.

Kim, N.Y., Ha, E., Moon, J.S., Lee, Y.H., and Choi, E.Y. (2020). Acute hyperglycemic crises with coronavirus disease-19: case reports. Diabetes Metab J 44, 349–353.

Klein, A.M., Mazutis, L., Akartuna, I., Tallapragada, N., Veres, A., Li, V., Peshkin, L., Weitz, D.A., and Kirschner, M.W. (2015). Droplet barcoding for single-cell transcriptomics applied to embryonic stem cells. Cell 161, 1187–1201.

Koch, M., Holmes, D., Bennet, N. (2020). COVID-19 and diabetes: a co-conspiracy? Lancet Diabetes Endocrinol 8, 801.

Kusmartseva, I., Wu, W., Syed, F., Van Der Heide, V., Jorgensen, M., Joseph, P., Tang, X., Candelario-Jalil, E., Yang, C., Nick, H., Harbert, J., Posgai, A., Lloyd, R., Cechin, S., Pugliese, A., Campbell-Thompson, M., Vander Heide, R., Evans-Molina, C., Homann, D., Atkinson, M. (2020). ACE2 and SARS-CoV-2 Expression in the Normal and COVID-19 Pancreas. Biorxiv.

Lan, J., Ge, J., Yu, J., Shan, S., Zhou, H., Fan, S., Zhang, Q., Shi, X., Wang, Q., Zhang, L., et al. (2020). Structure of the SARS-CoV-2 spike receptor-binding domain bound to the ACE2 receptor. Nature 581, 215–220.

Lee, I.T., Nakayama, T., Wu, C.T., Goltsev, Y., Jiang, S., Gall, P.A., Liao, C.K., Shih, L.C., Schurch, C.M., McIlwain, D.R., et al. (2020). Robust ACE2 protein expression localizes to the motile cilia of the respiratory tract epithelia and is not increased by ACE inhibitors or angiotensin receptor blockers. medRxiv.

Li, J., Wang, X., Chen, J., Zuo, X., Zhang, H., and Deng, A. (2020a). COVID-19 infection may cause ketosis and ketoacidosis. Diabetes Obes Metab.

Li, Y., Zhang, Z., Yang, L., Lian, X., Xie, Y., Li, S., Xin, S., Cao, P., and Lu, J. (2020b). The MERS-CoV receptor DPP4 as a candidate binding target of the SARS-CoV-2 spike. iScience 23, 101160.

Meireles, P.A., Bessa, F., Gaspar, P., Parreira, I., Silva, V.D., Mota, C., and Alvoeiro, L. (2020). Acalculous acute pancreatitis in a COVID-19 patient. Eur J Case Rep Intern Med 7, 001710.

Nair, G.G., Liu, J.S., Russ, H.A., Tran, S., Saxton, M.S., Chen, R., Juang, C., Li, M.L., Nguyen, V.Q., Giacometti, S., et al. (2019). Recapitulating endocrine cell clustering in culture promotes maturation of human stem-cell-derived beta cells. Nat Cell Biol 21, 263–274.

Picelli, S., Faridani, O.R., Bjorklund, A.K., Winberg, G., Sagasser, S., and Sandberg, R. (2014). Full-length RNA-seq from single cells using Smart-seq2. Nat Protoc 9, 171–181.

preprint: Fignani, D., Licata, G., Brusco, N., Nigi, L., Grieco, G., Marselli, L., Overbergh, L., Gysemans, C., Colli, M., Marchetti, P., Mathieu, C., Eizirik, D., Sebastiani, G., Dotta, F. (2020). SARS-CoV-2 receptor angiotensin I-converting enzyme type 2 is expressed in human pancreatic islet β-cells and is upregulated by inflammatory stress. BioRxiv.

Rabice, S.R., Altshuler, P.C., Bovet, C., Sullivan, C., and Gagnon, A.J. (2020). COVID-19 infection presenting as pancreatitis in a pregnant woman: A case report. Case Rep Womens Health 27, e00228.

Rafique, S., and Ahmed, F.W. (2020). A Case of Combined Diabetic Ketoacidosis and Hyperosmolar Hyperglycemic State in a Patient With COVID-19. Cureus 12, e8965.

Riddle, M.C., Buse, J.B., Franks, P.W., Knowler, W.C., Ratner, R.E., Selvin, E., Wexler, D.J., and Kahn, S.E. (2020). COVID-19 in people with diabetes: urgently needed lessons from early reports. Diabetes Care 43, 1378–1381.

Schreiber, B., Patel, A., and Verma, A. (2020). Shedding Light on COVID-19: ADAM17 the Missing Link? Am J Ther.

Segerstolpe, A., Palasantza, A., Eliasson, P., Andersson, E.M., Andreasson, A.C., Sun, X., Picelli, S., Sabirsh, A., Clausen, M., Bjursell, M.K., et al. (2016). Single-cell transcriptome profiling of human pancreatic islets in health and type 2 diabetes. Cell Metab 24, 593–607.

Seyedpour, S., Khodaei, B., Loghman, A.H., Seyedpour, N., Kisomi, M.F., Balibegloo, M., Nezamabadi, S.S., Gholami, B., Saghazadeh, A., and Rezaei, N. (2020). Targeted therapy strategies against SARS-CoV-2 cell entry mechanisms: A systematic review of in vitro and in vivo studies. J Cell Physiol.

Shang, J., Wan, Y., Luo, C., Ye, G., Geng, Q., Auerbach, A., and Li, F. (2020). Cell entry mechanisms of SARS-CoV-2. Proc Natl Acad Sci U S A 117, 11727–11734.

Stuart, T., Butler, A., Hoffman, P., Hafemeister, C., Papalexi, E., Mauck, W.M., 3rd, Hao, Y., Stoeckius, M., Smibert, P., and Satija, R. (2019). Comprehensive Integration of Single-Cell Data. Cell 177, 1888–1902 e1821.

Taneera, J., El-Huneidi, W., Hamad, M., Mohammed, A.K., Elaraby, E., and Hachim, M.Y. (2020). Expression Profile of SARS-CoV-2 Host Receptors in Human Pancreatic Islets Revealed Upregulation of ACE2 in Diabetic Donors. Biology (Basel) 9.

Teuwen, L.A., Geldhof, V., Pasut, A., and Carmeliet, P. (2020). COVID-19: the vasculature unleashed. Nat Rev Immunol 20, 389–391.

Uhlen, M., Fagerberg, L., Hallstrom, B.M., Lindskog, C., Oksvold, P., Mardinoglu, A., Sivertsson, A., Kampf, C., Sjostedt, E., Asplund, A., et al. (2015). Proteomics. Tissue-based map of the human proteome. Science 347, 1260419.

Unsworth, R., Wallace, S., Oliver, N.S., Yeung, S., Kshirsagar, A., Naidu, H., Kwong, R.M.W., Kumar, P., and Logan, K.M. (2020). New-Onset Type 1 Diabetes in Children During COVID-19: Multicenter Regional Findings in the U.K. Diabetes Care.

van den Brink, S.C., Sage, F., Vertesy, A., Spanjaard, B., Peterson-Maduro, J., Baron, C.S., Robin, C., and van Oudenaarden, A. (2017). Single-cell sequencing reveals dissociation-induced gene expression in tissue subpopulations. Nat Methods 14, 935–936.

Vankadari, N., and Wilce, J.A. (2020). Emerging WuHan (COVID-19) coronavirus: glycan shield and structure prediction of spike glycoprotein and its interaction with human CD26. Emerg Microbes Infect 9, 601–604.

Wang, F., Wang, H., Fan, J., Zhang, Y., Wang, H., and Zhao, Q. (2020). Pancreatic injury patterns in patients with coronavirus disease 19 pneumonia. Gastroenterology 159, 367–370.

Wang, S., Ma, P., Zhang, S., Song, S., Wang, Z., Ma, Y., Xu, J., Wu, F., Duan, L., Yin, Z., Luo, H., Xiong, N., Xu, M., Zeng, T., Jin, Y. (2020). Fasting blood glucose at admission is an independent predictor for 28-day mortality in patients with COVID-19 without previous diagnosis of diabetes: a multi-centre retrospective study. Diabetologia.

Wiersinga, W.J., Rhodes, A., Cheng, A.C., Peacock, S.J., and Prescott, H.C. (2020). Pathophysiology, Transmission, Diagnosis, and Treatment of Coronavirus Disease 2019 (COVID-19): A Review. JAMA.

Wright, J.J., Saunders, D.C., Dai, C., Poffenberger, G., Cairns, B., Serreze, D.V., Harlan, D.M., Bottino, R., Brissova, M., and Powers, A.C. (2020). Decreased pancreatic acinar cell number in type 1 diabetes. Diabetologia 63, 1418–1423.

Yang, J.K., Lin, S.S., Ji, X.J., and Guo, L.M. (2010). Binding of SARS coronavirus to its receptor damages islets and causes acute diabetes. Acta Diabetol 47, 193–199.

Yang, L., Han, Y., Nilsson-Payant, B.E., Gupta, V., Wang, P., Duan, X., Tang, X., Zhu, J., Zhao, Z., Jaffre, F., et al. (2020). A human pluripotent stem cell-based platform to study SARS-CoV-2 tropism and model virus infection in human cells and organoids. Cell Stem Cell 27, 125–136 e127.

Zang, R., Gomez Castro, M.F., McCune, B.T., Zeng, Q., Rothlauf, P.W., Sonnek, N.M., Liu, Z., Brulois, K.F., Wang, X., Greenberg, H.B., et al. (2020). TMPRSS2 and TMPRSS4 promote SARS-CoV-2 infection of human small intestinal enterocytes. Sci Immunol 5.

Zhou, F., Yu, T., Du, R., Fan, G., Liu, Y., Liu, Z., Xiang, J., Wang, Y., Song, B., Gu, X., et al. (2020). Clinical course and risk factors for mortality of adult inpatients with COVID-19 in Wuhan, China: a retrospective cohort study. Lancet 395, 1054–1062.

